# Temporal and protein-specific S-palmitoylation supports synaptic and neural network plasticity

**DOI:** 10.1101/2025.01.15.633145

**Authors:** Agata Pyty, Rabia Ijaz, Anna Buszka, Jacek Miłek, Iza Figiel, Patrycja Wardaszka, Matylda Roszkowska, Natalia Mierzwa, Adam Wojtas, Eli Kerstein, Remigiusz Serwa, Katarzyna Kalita, Rhonda Dzakpasu, Magdalena Dziembowska, Jakub Włodarczyk, Tomasz Wójtowicz

## Abstract

S-palmitoylation, a dynamic post-translational modification, has long been suggested to play a pivotal role in synaptic plasticity, learning, and memory. However, its precise impact on synaptic proteins and function remains unclear. In this study, we show that acute protein depalmitoylation in the hippocampus differentially affects short- and long-term synaptic plasticity, depending on synapse type. Strikingly, depalmitoylation also reprograms neuronal spiking timing following associative network activation. Our research identifies pre- and postsynaptic proteins dynamically regulated by S-palmitoylation during synaptic plasticity and suggests this modification occurs in isolated excitatory synapses. We also demonstrate that S-palmitoylation targets specific proteins within minutes and is not proteome-wide. These findings mark a significant advance in understanding how lipid modifications drive neural adaptability, memory, and learning.

## Introduction

A defining feature of the mammalian brain is its capacity to process and retain information within highly structured neuronal networks. Neuronal plasticity - the ability of neurons to adapt their connectivity, properties, and structure in response to activity - is fundamental to learning and memory. These processes involve the reorganization of existing synapses, modulation of synaptic efficacy, and adjustments to intrinsic neuronal excitability. Understanding the molecular mechanisms that govern these dynamic changes is central to advances in neuroscience, neuropharmacology, and medical research.

Post-translational modifications (PTMs) add a critical layer of regulation to the proteins driving neuronal plasticity. By modulating protein function, localization, stability, and interactions, PTMs enable the rapid and precise tuning of cellular processes necessary for synaptic remodeling and network adaptation. Among PTMs, lipid modifications have emerged as key regulators of neuronal function. These modifications, including N-myristoylation, N-acylation, O-acylation, and S-acylation, alter the hydrophobicity of proteins, driving their association with membranes or specific membrane microdomains (1). S-acylation is reversible by deacylation and can therefore act as a dynamic on/off switch, similar to phosphorylation, acetylation, and ubiquitynation. Since the 16-carbon-long palmitic acid is the most frequently added moiety, S-acylation has often been referred to as S-palmitoylation (S-PALM), or simply palmitoylation, even when the specific lipid has not been identified (1). Over the past two decades, S-PALM has emerged as a crucial regulator of protein function. S-PALM was found to alter the conformation or structure of a protein, intracellular trafficking and membrane localization, protein-protein interactions and signal transduction (2,3). This process is mediated by palmitoyl acyltransferase enzymes (PATs) and reversed by thioesterases, (reviewed in (4–6)) and may be potentially important in the context of activity-dependent modification of synapse function. Protein palmitoylation is not only thought to contribute to healthy physiological processes such as learning and memory, but may play a role in neurodegenerative diseases such as Alzheimer’s disease (AD), Parkinson’s disease (PD), Huntington’s disease (HD), amyotrophic lateral sclerosis (ALS) and other neuropsychiatric disorders (4,7).

In the nervous tissue, S-PALM was extensively studied in major synaptic proteins, such as the scaffolding postsynaptic density protein 95 (PSD-95) and gephyrin, as well as ionotropic receptors (α-amino-3-hydroxy-5-methyl-4-isoxazolepropionic acid receptors [AMPARs], N-methyl-D-aspartate receptors [NMDARs], and γ-aminobutyric acid receptors [GABARs]) that mediate the majority of fast excitatory and inhibitory synaptic transmission in the brain (4–6,8). Emerging evidence suggests that S-PALM may serve as an additional mechanism to orchestrate synaptic protein distribution, thereby supporting synaptic plasticity, learning and memory (4). For instance, manipulation of several PATs genes such as ZDHHC3, ZDHHC7, ZDHHC9, or ZDHHC17, was reported to affect basal excitatory and/or inhibitory synaptic transmission (4), while knockout of the gene coding depalmitoylating enzyme PPT1 was shown to affect inhibitory synaptic transmission and presynaptic release (9,10). Moreover, knockouts of PATs including ZDHHC2, ZDHHC5, ZDHHC9, and ZDHHC17, as well as PPT1, were linked with impairments in spatial or fear learning (9,11–13) and pharmacological inhibition of palmitoylation impairs memory acquisition and consolidation in the hippocampus (14). These findings indicate that palmitoylation may regulate the excitatory-inhibitory balance in neuronal networks and play a vital role in normal brain function, particularly in neuronal plasticity. A recent study revealed a list of > 60 synaptic proteins that could undergo changes in S-PALM 1h after the formation of a context-dependent fear memory in mice (15), suggesting that this process is highly protein-specific. However, the mechanisms underlying protein selectivity in neuronal S-palmitoylation, the precise temporal dynamics of this modification, and the signals that trigger its regulation remain poorly understood. Unraveling these aspects is critically important, as dysregulated S-palmitoylation has been implicated in numerous neuropsychiatric conditions.

Functional studies conducted both *in vitro* and *in vivo* have confirmed that memory traces can be encoded through activity-dependent modifications of synapses. A defining feature of such synaptic plasticity is long-term potentiation (LTP) or long-term depression (LTD), which are elicited by patterned stimulation of afferent fibers at high and low frequencies, respectively (16). The most widely studied form of synaptic plasticity is LTP, which leads to long-lasting strengthening of synapses in response to enhanced synapse activity (17) and is considered the biological substrate for learning and memory (18). Synaptic potentiation can occur through pre- and / or postsynaptic mechanisms and may involve NMDAR subtype of glutamate receptors (19), although some forms of LTP can be expressed independently of NMDAR activity. Regardless of the specific mechanism, most glutamatergic synapses require an initial rise in cytosolic Ca^2+^ concentrations for LTP induction. This increase is mediated via NMDARs, voltage-gated calcium channels (VGCCs), calcium-permeable AMPA receptors (CP-AMPARs), or release from intracellular calcium stores (17). The early phase of LTP (E-LTP) primarily depends on protein trafficking to support synaptic potentiation. In contrast, the late phase of LTP (L-LTP) involves longer-lasting, transcription-dependent changes that stabilize synaptic modifications, resulting in sustained synaptic enhancement (20). Additional factors contribute to LTP induction and maintenance, such as brain-derived neurotrophic factor (BDNF), which facilitates synaptic potentiation by activating the TrkB receptor tyrosine kinase (21).

In this study, we combined electrophysiology, biochemistry, proteomics, and imaging to explore the role of protein palmitoylation in synaptic plasticity and behavior of neuronal networks. By inducing plasticity in various models, we mapped the temporal dynamics of S-palmitoylation, assessed its impact on excitatory synapse function, and identified proteins dynamically modified following increased neuronal activity. Our findings reveal that S-PALM supports the plasticity of excitatory transmission in the hippocampus in a synapse-dependent manner and may modulate neuronal spiking and information flow within neural networks. At the molecular level, we observed that the induction of synaptic plasticity prompts transient and protein-specific palmitoylation.

## Results

### The induction of neuronal plasticity in vitro regulates the palmitoylation of synaptophysin and neurochondrin

Understanding neuronal assemblies is essential for uncovering the mechanisms underlying brain coding, learning, and memory. To explore protein palmitoylation’s role in activity-dependent neuronal spiking and network properties, at 14 days *in vitro* (DIV) primary rat hippocampal neurons were treated with picrotoxin, forskolin, and rolipram for 20 or 60 minutes to induce long-term synaptic plasticity. This cocktail is known to increase cAMP level and network activity leading to a tetanic-like stimulation in bulk and induction of functional and structural neuronal plasticity (22,23) (Fig 1 A). This method of neuronal stimulation is referred to as chemically induced long-term potentiation (cLTP). We demonstrated that 1 hour after cLTP induction, there is a significant upregulation of phosphorylated extracellular signal-regulated kinase (pERK), a molecular marker of LTP. (Fig 1 C) (24). We next studied protein palmitoylation and subjected homogenized neuronal cultures to the acyl-biotin exchange (ABE) method (25) and quantified total protein palmitoylation levels as well as that of individual proteins (see methods for details). We observed that cLTP induction did not affect global protein palmitoylation levels (Fig. 1B) when normalized to total protein levels (stain-free technology, not shown). However, it significantly altered the palmitoylation status of specific presynaptic and postsynaptic proteins (Fig 1 D-F). Specifically, we examined three categories of proteins: presynaptic (synaptophysin, vesicle-associated membrane protein 2 [VAMP2], synaptosome-associated protein 25 [SNAP25]), postsynaptic (postsynaptic density protein 95 [PSD-95], glutamate ionotropic receptor AMPA type subunit 1 [GluR1], neurochondrin), and inter-synaptic (neural cell adhesion molecule [NCAM]). At 1 hour post-cLTP induction, the palmitoylation levels of synaptophysin and neurochondrin were significantly decreased (Fig 1 D–F). In contrast, in the same samples, the palmitoylation level of PSD-95 was significantly increased, consistent with previous findings (26,27) while most other proteins we analyzed did not exhibit significant changes (Fig 1 D–F). To further explore the time-dependence of these changes, we repeated the experiment with a shorter time window of 20 minutes. Unlike the 1-hour treatment, at 20 minutes post-cLTP induction, the palmitoylation level of synaptophysin increased, that of NCAM decreased, and GluR1 remained unaltered (Fig S1). Overall, these results demonstrate that the induction of neuronal plasticity via cLTP in primary neuronal cultures leads to time-dependent and protein-specific changes in protein palmitoylation. We indicate synaptophysin, neurochondrin, and NCAM as examples of dynamic regulation by palmitoylation.

**Fig 1.**
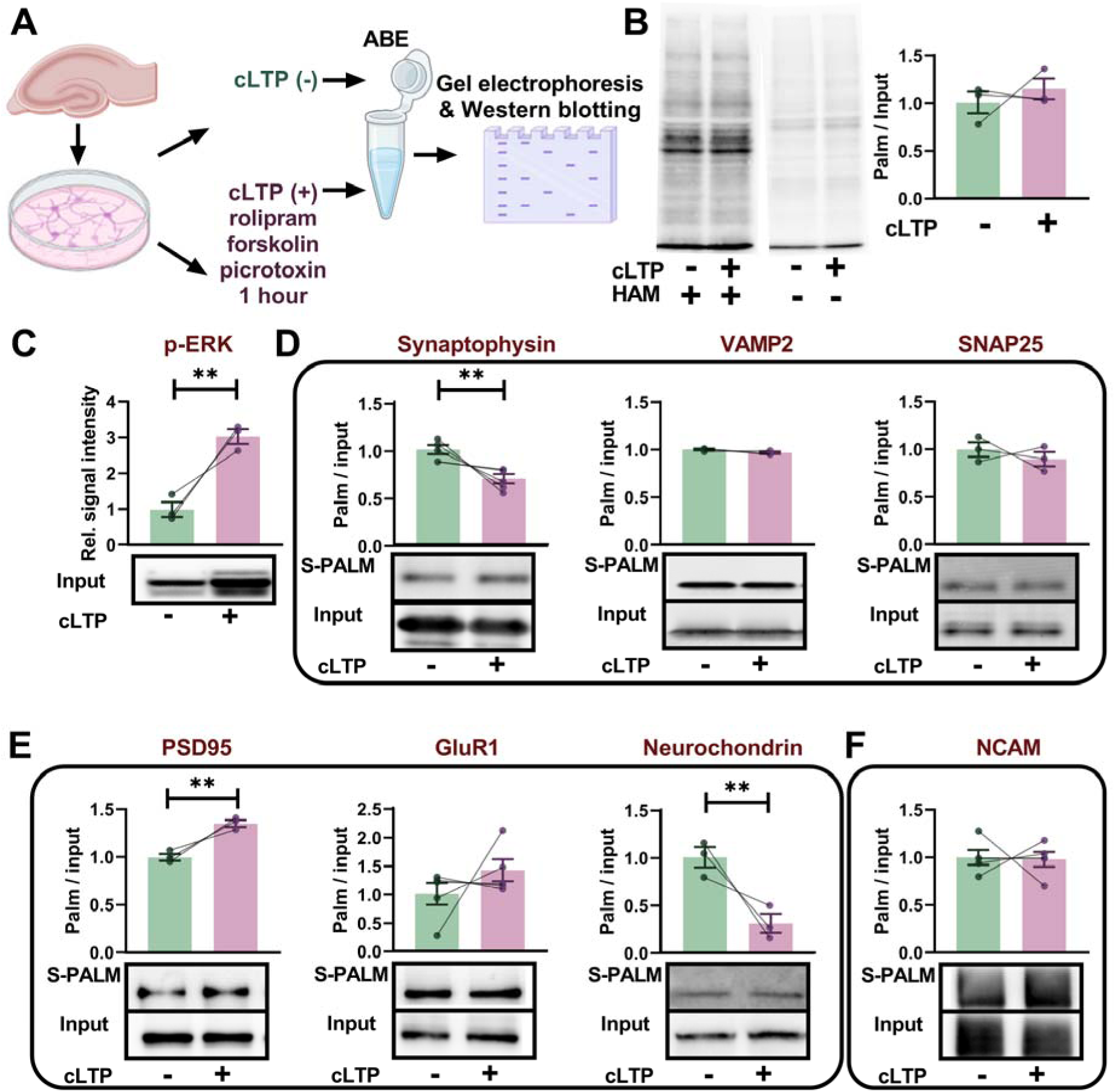
Induction of synaptic plasticity in neuronal cultures regulates palmitoylation of synaptic proteins. (A) Primary rat hippocampal neurons were cultured for 14 days and treated with cLTP cocktail (cLTP +) or solvent (cLTP -). Culture homogenates were collected 1 hour after treatment, subsequentl subjected to ABE and S-PALM and input fractions were immunoblotted for target proteins, as indicated. (B) Western blot of global palmitoylation at 1 hour post cLTP. HAM stands for hydroxylamine. There was no shift in global protein palmitoylation (*n =* 3 cultures, *p =* 0.40, unpaired Student’s t-test). (C) Quantification of the phosphorylated extracellular regulated kinase (pERK) protein expression in cultur homogenates. cLTP resulted in a significant increase in the expression of pERK protein (*n =* 3 cultures, *p =* 0.002, unpaired Student’s t-test). (D) Western blot and quantification of the ABE assay described in (A) performed on selected presynaptic proteins. The levels of palmitoylated synaptophysin and neurochondrin significantly decreased post cLTP (*n =* 5 and 3 cultures respectively, *p =* 0.002 and *p =* 0.009, respectively, unpaired Student’s t-test). (E) Western blot and quantification of the ABE assay described in (A) performed on selected postsynaptic proteins. The levels of palmitoylated PSD-95 increased significantly post cLTP (*n =* 3 cultures, *p =* 0.002, unpaired Student’s t-test). (F) Western blot and quantification of the ABE assay described in (A) performed on an exemplary bipolar adhesion molecule NCAM. Data are mean ± SEM from *n =* 3 to 5 cultures. \*\**p <* 0.01.

### Protein deacylation alters the temporal organization of neuronal spiking following periods of heightened network activity

To manipulate protein palmitoylation, at 13 DIV we treated neurons overnight with deacylation agent N-(tert-Butyl)hydroxylamine (NtBuHA), a hydroxylamine derivative (28). As shown in Fig S2 (A-B), the chemiluminescent signal corresponding to total proteome palmitoylation in culture homogenates was significantly reduced following overnight treatment with NtBuHA (1 mM). Moreover, NtBuHA at 1 mM, as well as 2-bromopalmitate (2-BP) (100 µM) an irreversible and nonselective inhibitor of palmitoyl acyltransferases (4,29), did not induce cellular toxicity, as indicated by the absence of increased lactate dehydrogenase (LDH) release (Fig S2 C). To further evaluate the efficacy of NtBuHA as a deacylation agent, total proteome palmitoylation in neuronal cultures was assessed using a click chemistry reaction with Oregon Green 488 dye (30). Primary rat hippocampal cultures at 13 DIV were treated overnight with exogenous alkyne-palmitate to facilitate its incorporation into cellular proteins. Some cultures were additionally exposed to 1 mM NtBuHA. As shown in Fig S2 (D-E), NtBuHA significantly reduced fluorescence associated with labeled exogenous palmitate to the level of negative control (cultures treated only with the fluorophore), further confirming its effectiveness in globally deacylating neuronal proteins. Based on these findings, we utilized water-soluble NtBuHA (1 mM) in subsequent experiments.

We investigated how protein depalmitoylation impacts long-term, activity-dependent changes of neural responses in cultured rat primary hippocampal neurons. Cells were cultured for 14 days on multielectrode arrays (MEA), allowing for extracellular and non-invasive recordings of action potentials in a network of neurons from up to 60 sites (Fig 2 A-B). These multisite recordings facilitate the capture of spiking changes at the population level while allowing for the manipulation of neuronal activity via electrical stimulation. Under basal conditions, control (CTR) MEA exhibited 39 ± 4.6 active electrodes, an average spike frequency of 2.6 ±0.49 Hz, and a mean burst duration (≥ 5 spikes) of 132 ± 19 ms (n = 5 MEA, Fig S3). To induce network plasticity, we first attempted medium exchange combined with the addition of a cLTP cocktail; however, this approach was ineffective (see Discussion). Consequently, we applied paired associative activity at two electrodes within the network of hippocampal neurons for 2 minutes (AT-IN protocol, Fig 2 C, see Methods (29)) and subsequently monitored network activity for 1 hour. Such a period of enhanced neuronal activity or the presence of NtBuHA did not affect the number of active electrodes, number of bursts or the average spike frequency (Fig S3). However, electrical stimulation had a significant impact on the temporal organization of spiking (Fig 2 D). The Fano Factor reflects the variability in spike occurrence relative to the average number of spikes. In the CTR group, the Fano Factor significantly decreased over time suggesting that the network became more organized with more regular or rhythmic firing. In contrast, following deacylation with NtBuHA, an inverse effect was observed (Fig 2 D). Altogether, protein deacylation had no effect on the mean rate or temporal distribution of action potentials in the neuronal network, but it impaired changes in the temporal organization of neuronal spiking following a period of enhanced network activity.

**Fig 2.**
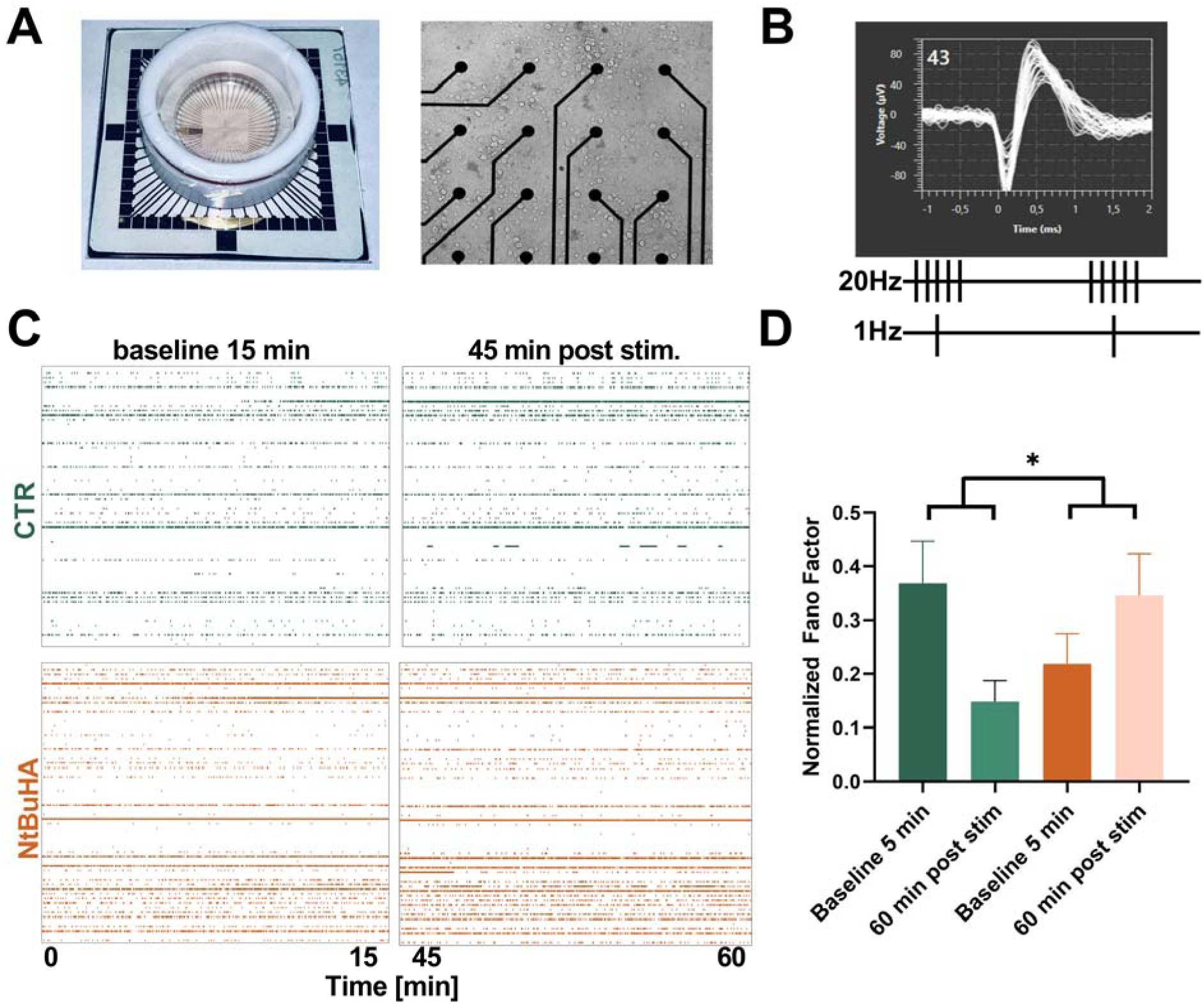
Protein deacylation modulates the temporal organization of neuronal spiking following a period of enhanced network activity. (A) Primary rat hippocampal neurons were cultured for 14 days on multielectrode arrays (MEA), allowing recordings of action potentials in a network of neurons from up to 60 sites. MEAs were covered with fluorinated ethylene-propylene membranes to maintain medium osmolarity and the day before recordings received either water (0.01%) or deacylation agent NtBuHA (1 mM, overnight). (B) Representative waveforms of extracellularly recorded action potentials at 14 DIV together with a scheme of pattern of electrical stimulation. Network plasticity was analyzed up to 30 minutes before and up to 60 minutes after electrical stimulation of two non-adjacent electrodes on the MEA (20 Hz bursts paired with 1 Hz in-phase activity for a total of 120 seconds). (C) Exemplary raster plots show spiking activity over a 15-minute period before and after associative electrical stimulation. Each row corresponds to an electrode, and each dot represents a single action potential. (D) Quantification of spikes recorded in control cultures (green) and cultures treated overnight with NtBuHA (orange) before and after inducing network plasticity through associative activity (stim.). The normalized Fano Factor was analyzed before and after stimulation. A significant interaction between treatment and stimulation was observed (*F*(1,8) = 7.421, *p* = 0.026, two-way ANOVA) but no significant main effects were found. Data are shown as mean ± SEM. *n =* 5 MEAs per group. Asterisk indicates statistical significance (\**p <* 0.05).

### NMDAR-dependent synaptic potentiation in the hippocampus results in rapid and protein-specific palmitoylation

We next analyzed protein palmitoylation in acute hippocampal slices following induction of LTP. To induce NMDAR-dependent LTP, we incubated hippocampal slices in Mg^2+^-free aCSF with addition of glycine (600 µM) for 10 minutes and subsequently maintained the slices in the same aCSF without glycine for 20 minutes (glycine LTP, gLTP) (31). To confirm the efficacy of this protocol, we performed recordings of evoked field excitatory postsynaptic potentials (fEPSPs) in response to Shaffer collaterals stimulation. We recently discovered that the molecular mechanisms of synaptic plasticity and the protein involvement in neighboring excitatory connections on basal and apical dendrites in CA1 hippocampal region differ significantly (32), similarly to synapses in pyramidal neurons of CA3 region (33). Therefore, we looked at synaptic signals in *stratum oriens* (SR) and *stratum radiatum* (SO) (a hub to excitatory synapses located in basal or apical dendrites of pyramidal neurons, respectively). As shown in the Fig S4 (A-D), the application of glycine at time 0 for 10 minutes resulted in significant potentiation of fEPSP amplitudes in both regions. To confirm whether gLTP is NMDAR-dependent, we repeated recordings in SR with the addition of the NMDAR competitive antagonist APV (50 μM) to the glycine solution. In the presence of APV, the glycine effect was not observed (Fig S4 A). In addition, detected fEPSP amplitudes were significantly increased in the presence of glycine in both SR and SO but not when glycine was co-applied with APV (Fig S4 B-D). We also found in hippocampal slice homogenates that gLTP resulted in a significant upregulation of activity-regulated cytoskeleton-associated protein (Arc), a molecular marker of LTP (Fig 3 C) (34). Altogether, these results confirm that glycine-induced synaptic plasticity in acute hippocampal slices occurs in an NMDAR-dependent way and provide a reliable model to study protein-specific palmitoylation following synaptic potentiation.

**Fig 3.**
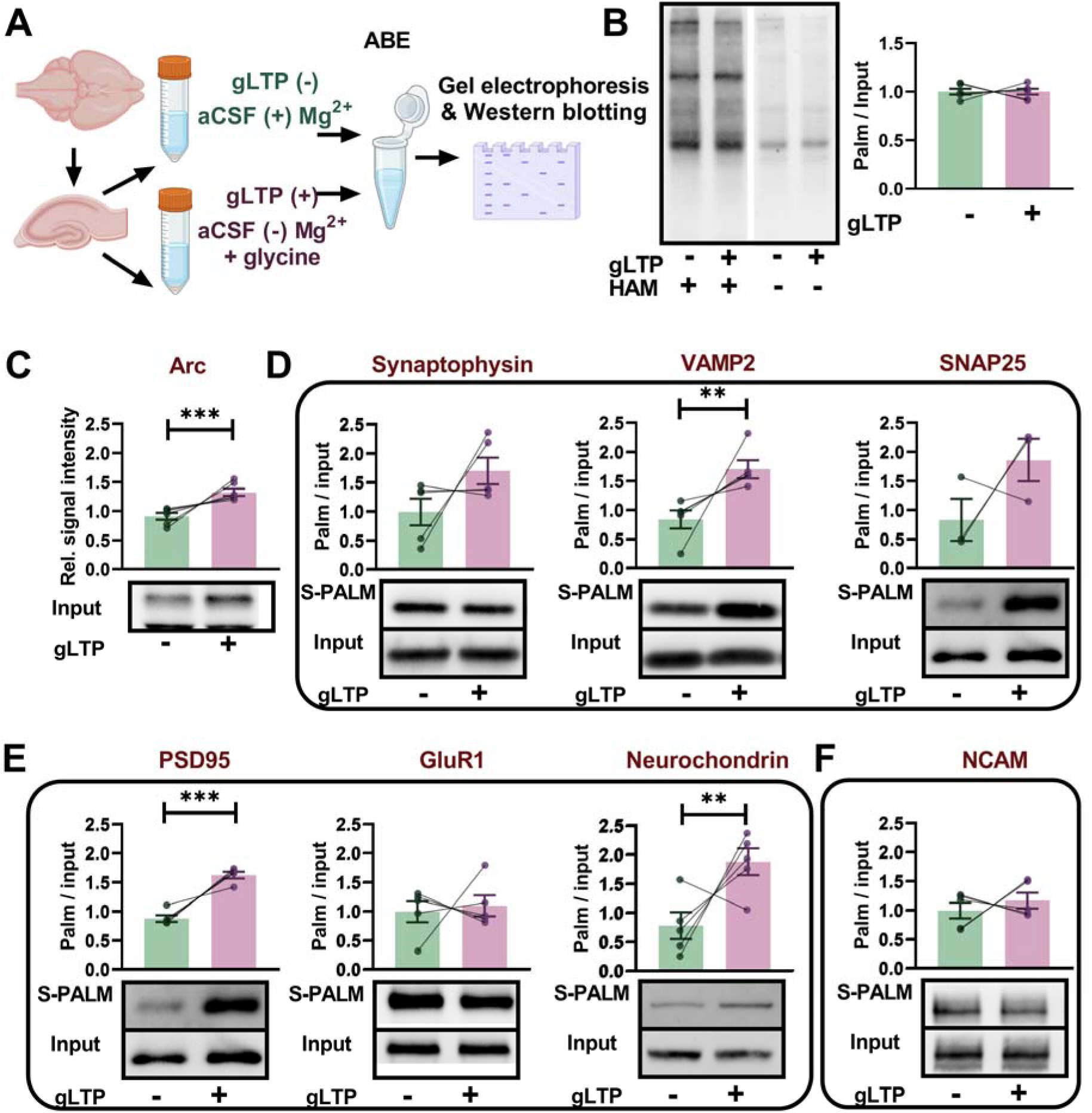
Glycine-induced LTP (gLTP) in hippocampal slices induces the differential palmitoylation of proteins. (A) Schematic illustration of the NMDAR-dependent glycine induced LTP assay in hippocampal slices to investigate activity-induced changes in neuronal protein palmitoylation. Tissue homogenates were collected 20 min after treatment with either mock gLTP (−) or gLTP (+). Both S-PALM and input fractions were immunoblotted for target proteins, as indicated. (B) Western blot of global palmitoylation at 20 minutes post glycine induced cLTP. HAM stands for hydroxylamine. Glycine LTP did not affect global protein palmitoylation levels (*n =* 6 animals, *p =* 0.99, unpaired Student’s t-test). (C) Quantification of the Arc protein expression in tissue homogenates. Glycine induced LTP resulted in significant increase in the expression of Arc protein (*n =* 6 animals, *p <* 0.006, unpaired Student’s t-test). (D) Western blot and quantification of the ABE assay described in (A) performed on selected presynaptic proteins. The levels of palmitoylated VAMP2 increased significantly post cLTP (*n =* 5 animals, *p =* 0.004, unpaired Student’s t-test). (E) Western blot and quantification of the ABE assay described in (A) performed on selected postsynaptic proteins. The levels of palmitoylated PSD-95 and neurochondrin increased significantly post cLTP (*n =* 5 animals, *p <* 0.0001 and *p =* 0.009, respectively, unpaired Student’s t-test). (F) Western blot and quantification of the ABE assay described in (A) performed on an exemplary bipolar adhesion molecule NCAM. Data are mean ± SEM. \*\**p <* 0.01, \*\*\**p <* 0.001.

We next investigated how gLTP affected palmitoylation of synaptic proteins in slice homogenates subjected to the ABE method (25) (see methods for details). We found that gLTP did not affect global protein palmitoylation chemiluminescent signal (Fig 3 B) when normalized to total protein signal (stain-free technology, not shown). However, it significantly altered the palmitoylation level of certain presynaptic or postsynaptic proteins. In contrast to neuronal cultures, most proteins exhibited hyperpalmitoylation. In particular, VAMP2, neurochondrin and PSD-95 palmitoylation levels were significantly increased (Fig. 3 D-E). Palmitoylation levels of synaptophysin and SNAP25 showed an upward trend, although these changes did not reach statistical significance. Altogether, these results demonstrate that NMDAR-dependent synaptic plasticity in hippocampal slices induces rapid and protein-specific upregulation of palmitoylation, highlighting the dynamic regulation of lipid modifications in synaptic function.

### Protein depalmitoylation differentially affects long-term synaptic potentiation in hippocampal excitatory synapses

Next, we studied whether involvement of protein palmitoylation in synaptic plasticity depends on the type of excitatory synapse. Before electrophysiological recordings, slices were bathed in NtBuHA (1 mM) for 2 hours. Incubation of hippocampal slices with NtBuHA significantly downregulated protein palmitoylation in homogenates, as shown by the ABE method (Fig S5 A-B). fEPSPs were then evoked in SO or SR in response to Schaffer collaterals stimulation (see Methods for details). As shown in Fig. 4 (B, G), fEPSPs amplitudes recorded in response to monotonically increasing stimuli (input-output curves) were significantly increased in the presence of NtBuHA in SR but not in SO. Next, we studied short-term synaptic plasticity by stimulating afferents with two pulses with 50 ms inter-event interval. In control slices, paired-pulse facilitation (PPF) ratio was 1.67 ± 0.12 (*n =* 19 slices) in SR and 1.48 ± 0.04 in SO (*n =* 16 slices). We observed a significant reduction of paired-pulse facilitation ratio in SR in the presence of NtBuHA but not in SO (Fig. 4 C, H). Altogether, in the presence of the deacylating agent NtBuHA, basal synaptic transmission and short-term facilitation were affected preferentially in SR but not in SO.

**Fig 4.**
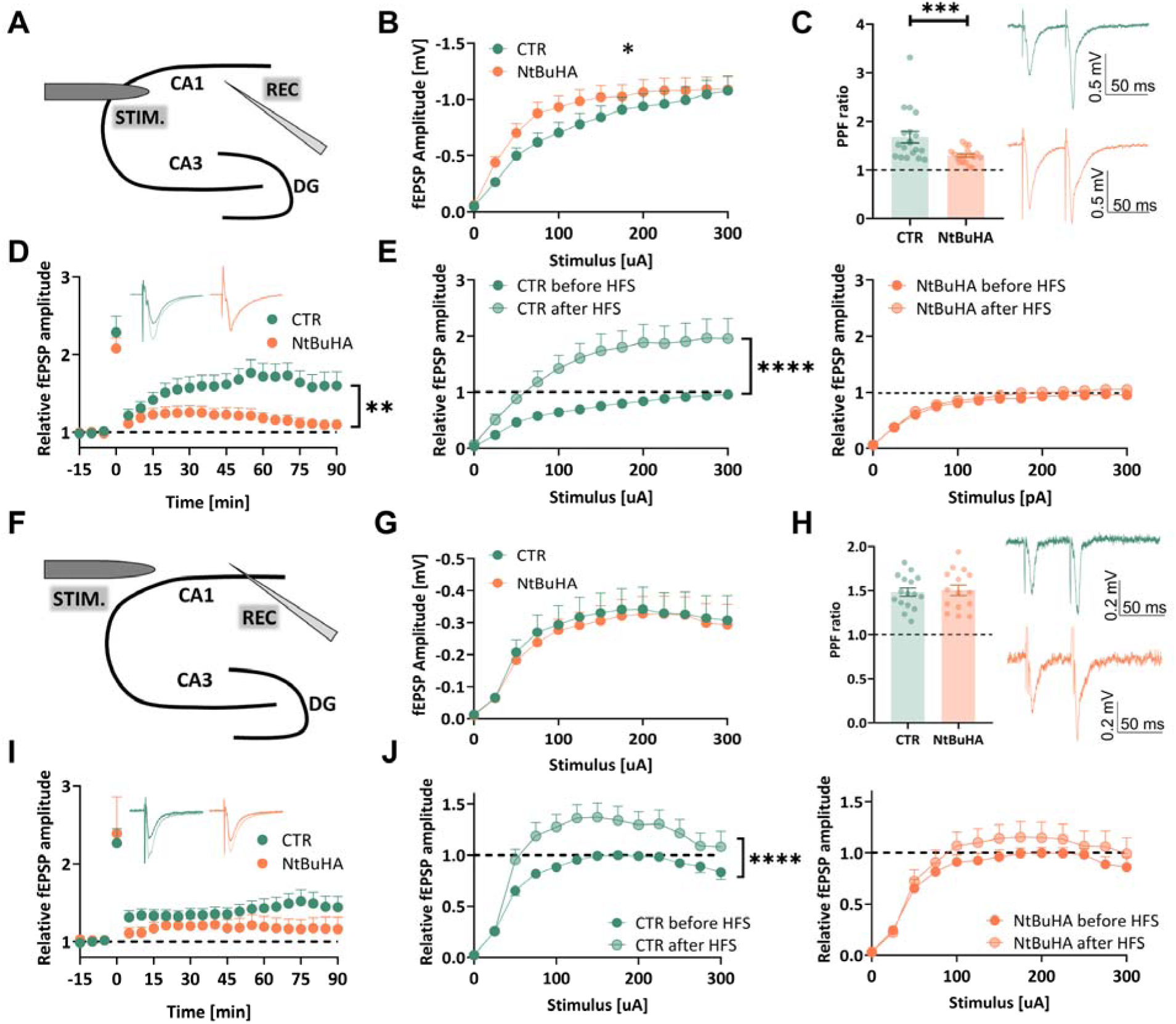
Protein depalmitoylation differentially affects long-term synaptic potentiation in hippocampal excitatory synapses. (A) Schematic representation of fEPSP recordings in SR excitatory synapses in response to Schaeffer collaterals stimulation (A-E). (B) Statistical analysis of fEPSP amplitudes recorded in response to extracellular stimulation in the control slices and those incubated with depalmitoylating agent NtBuHA. Note that in the presence of the drug, fEPSP amplitudes were increased (*n =* at least 17 slices per group, *F*(12,396) = 2.044, *p =* 0.019, two-way ANOVA, treatment × stimulus intensity). (C) Statistical evaluation of fEPSP recordings in response to paired-pulse stimulation (50ms inter-stimulus-interval). It is noteworthy that in the presence of the drug, the index is significantly decreased (*p =* 0.0052, unpaired-t-test). Insets show exemplary recordings with matching colors. (D) Exemplary traces of fEPSPs and the time course of fEPSPs recorded before and after LTP induction with high frequency stimulation (hfsLTP, 4 × 100 Hz, applied at time = 0 min.). In the presence of NtBuHA, the magnitude of hfsLTP at 90 minutes was significantly decreased (*n =* 15 and *n =* 16 for CTR and NtBuHA groups, respectively, *p =* 0.013, unpaired t-test). (E) Statistics of fEPSP responses to monotonically increased stimuli before (full color) and 90 minutes after hfsLTP induction (transparent color). In control slices, hfsLTP resulted in significant upregulation of fEPSP amplitudes in response to a wide range of stimuli (left panel, *F*(12, 408) = 6.449, *p <* 0.0001, two-way ANOVA, treatment × stimulus intensity) and this was not observed in slices treated with NtBuHA (right panel, *F*(12, 384) = 1.005, *p =* 0.44, two-way ANOVA). (F) Schematic presentation of fEPSP recordings in SO excitatory synapses in response to Schaeffer collaterals stimulation (G-J). (G) Statistical analysis of fEPSP amplitudes recorded in response to extracellular stimulation. There was no difference between the control slices and those incubated with the depalmitoylating agent NtBuHA (*n =* at least 15 slices per group, *F*(12,348) = 0.07, p > 0.99, two-way ANOVA, treatment × stimulus intensity). (H) Statistics of fEPSP recordings in response to paired-pulse stimulation (50ms inter-stimulus-interval) indicating no difference between investigated groups (*p =* 0.78, unpaired-t-test). Insets show exemplary recordings with matching colours. (I) Exemplary traces of fEPSPs and the time course of fEPSPs recorded before and after hfsLTP induction with high-frequency stimulation (4 × 100 Hz, applied at time = 0 min.). In the presence of NtBuHA, the magnitude of hfsLTP at 90 minutes was not significantly different from controls (*n =* 15 and *n =* 16 for CTR and NtBuHA groups, respectively, *p =* 0.15, unpaired t-test). (J) Statistics of fEPSP responses to monotonically increased stimuli before (full color) and 90 minutes after hfsLTP induction (transparent color). In control slices, hfsLTP resulted in significant upregulation of fEPSP amplitudes in response to a wide range of stimuli (left panel, *F*(12, 360) = 2.854, *p =* 0.0009, two-way ANOVA, treatment × stimulus intensity), which was not observed in slices treated with NtBuHA (right panel, *F*(12, 336) = 0.417, *p =* 0.95, two-way ANOVA, treatment × stimulus intensity). Asterisks indicate statistical significance: * *p <* 0.05, ** *p <* 0.01, *** *p <* 0.01, **** *p <* 0.0001.

We next studied the impact of protein deacylation on long-term synaptic plasticity induced with high-frequency stimulation (4 × 100 Hz) of Schaffer collaterals (hfsLTP). We found that in control (CTR) slices, the relative fEPSP amplitude 90 minutes post hfsLTP induction was 1.60 ± 0.18 (*n =* 16 slices) in SR and 1.44 ± 0.12 (*n =* 16 slices) in SO. hfsLTP was significantly impaired in the presence of NtBuHA in SR (*n =* 16 slices, *p =* 0.013, unpaired t-test) but not in SO (*n =* 15 slices, *p =* 0.15, unpaired t-test, Fig. 4. D, I). Induction of hfsLTP resulted in a significant leftward shift in the input-output function in control slices (F(_12, 360_) = 6.749, *p <* 0.0001, *n =* 16 slices, two-way ANOVA) and this shift was not observed in the presence of NtBuHA (F(_12, 384_) = 1.005, p=0.44, two-way ANOVA). Similarly, in SO, hfsLTP induction resulted in a significant leftward shift in the input-output function in control slices (F(_12, 360_) = 2.854, *p =* 0.0009, two-way ANOVA) but this shift was not observed in NtBuHA group (F(_12, 336_) = 0.41, *p =* 0.95, two-way ANOVA) (Fig. 4 E, J). In summary, protein depalmitoylation differentially affects synaptic plasticity in SR and SO. While basal transmission, the time course of LTP, and short-term facilitation were preferentially affected in SR, both synaptic regions exhibited impaired changes in the magnitude of synaptic responses to a wide range of stimuli following LTP.

### Excitatory synapses contain palmitoylation machinery responsive to external stimuli

Most PATs are transmembrane enzymes located in the Golgi apparatus and the endoplasmic reticulum (4). In dendrites and spines several members of the PATs family such as ZDHHC2, ZDHHC5, ZDHHC8, and ZDHHC17 have been suggested to regulate the location and function of multiple signaling proteins (6,27). Given the presence of PATs in dendrites and spines, we investigated whether isolated synaptoneurosomes contain active palmitoylation machinery capable of responding dynamically to cLTP stimuli. Synaptoneurosomes contain both presynaptic (synaptosome) and postsynaptic (neurosome) vesicularized components, and are widely used to study synaptic structure and function (35). It was reported that isolated synaptoneurosomes can undergo LTP-like plasticity in response to stimuli that mimic synaptic NMDAR activation (36). In this protocol, KCl-evoked release of endogenous glutamate from presynaptic terminals, in the presence of the NMDAR co-agonist glycine, leads to a long-lasting increase in surface AMPAR levels (Fig. 5 A).

**Fig 5.**
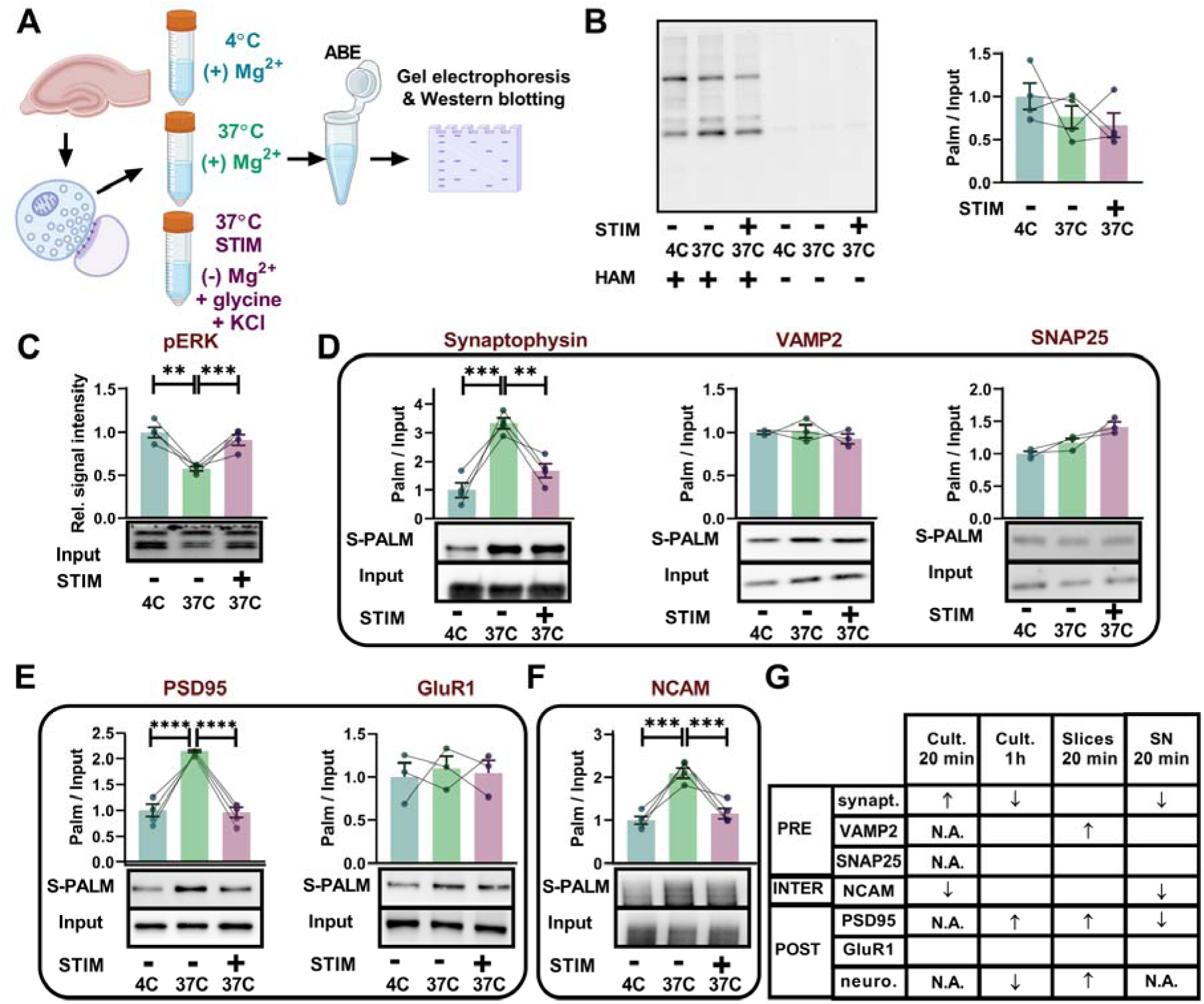
Synaptoneurosomes contain a palmitoylation machinery responsive to external stimuli. (A) Schematic illustration of the NMDAR-dependent glycine and KCl induced stimulation assay (STIM) in live synaptoneurosomes from hippocampi of young rats to investigate activity-induced changes in synaptic protein palmitoylation. Synaptoneurosomes were either kept on ice (4C) or treated with either sham (−) or STIM (+) for 20 minutes. Both S-PALM and input fractions were immunoblotted for target proteins, as indicated. (B) Western blot of global palmitoylation indicates no global change in protein palmitoylation of synaptoneurosomes post STIM (*n =* 4 animals, *p =* 0.27, *F*(2, 9) = 1.486, one-way ANOVA). (C) Quantification of the pERK protein expression in synaptoneurosomes. STIM resulted in an increase in ERK phosphorylation compared to mock solution (-) (*n =* 4 animals, *F*(2, 9) = 17.27, *p <* 0.001, one-way ANOVA). (D) Western blot and quantification of the ABE assay described in (A) performed on selected presynaptic proteins. The levels of palmitoylated synaptophysin decreased significantly post STIM (*n =* 4 animals, *F*(2, 9) = 27.49, *p* < 0.001, one-way ANOVA). (E) Western blot and quantification of the ABE assay described in (A) performed on selected postsynaptic proteins. Th level of palmitoylated PSD-95 decreased significantly post STIM (*n =* 4 animals, *p <* 0.001, *F*(2, 9) = 53.31, one-way ANOVA). (F) Western blot and quantification of the ABE assay described in (A) performed on exemplary bipolar adhesion molecule NCAM. The level of palmitoylated NCAM decreased significantly post STIM (n= 4 animals, *F*(2, 9) = 27.79, *p <* 0.001, one-way ANOVA). (G) Summary of results obtained in experimental models of synaptic plasticity: neuronal cultures, hippocampal slices and synaptoneurosomes. Arrows indicate the change in palmitoylation levels relative to the control for each protein. N.A. indicates not applicable (not studied). SN - synaptoneurosomes. Empty cell indicates palmitoylation level did not change significantly. Data are mean ± SEM. \*\**p <* 0.01, \*\*\**p <* 0.001, \*\*\*\**p <* 0.0001).

Consequently, we prepared synaptoneurosomes from rat hippocampi in ice-cold solutions (see Methods), warmed them at 37°C for 10 minutes, and subsequently treated them with a stimulating solution (STIM) containing glycine and 20 mM KCl for 20 minutes (36) (see Methods for details). Additionally, for each replicate part of the material was left on ice (4°C) throughout the experiment to check the effect of temperature on protein stability and palmitoylation. The STIM protocol resulted in a significant upregulation of the level of phosphorylated extracellular signal regulated kinase (pERK), a molecular marker of LTP (Fig. 5 C) when compared to the control sample 37°C (24). In contrast, control 37°C samples had significantly less pERK compared to 4°C samples (Fig 5C). Having confirmed that synaptoneurosomes are viable and responsive to extracellular stimuli (temperature and STIM), we used the ABE method on synaptoneurosomal fractions to analyze palmitoylation levels of proteins. We found that stimulation did not affect global protein palmitoylation in synaptoneurosomes (Fig 5 B). However, it significantly altered palmitoylation levels of selected presynaptic or postsynaptic proteins described in Fig. 1. We found that in contrast to neuronal cultures and slices, most proteins exhibited depalmitoylation following stimulation. In particular, synaptophysin, PSD-95 and NCAM palmitoylation levels were significantly decreased (Fig. 5 D-F). A summary of the changes in palmitoylation for all the proteins investigated here (cultures, slices and synaptoneurosomes) is shown in Fig 5 G. Altogether, the stimulation of synaptoneurosomes with glycine and KCl led to rapid and protein-specific downregulation of palmitoylation.

To further investigate the palmitoylome of synaptoneurosomes, we performed a mass spectrometry-based proteomic analysis (see Methods for details; Data S1). This analysis of the baseline proteome identified 4,469 proteins, including two palmitoyltransferases (ZDHHC17 and ZDHHC5) and six thioesterases (e.g., Lypla1, Lypla2, Ppt1, and Abhd17a-c) (Data S1, Fig 6). The expression level of 90 proteins (2%) was altered following STIM. Therefore, all the palmitoyl-proteins were normalized to their input. Among the 4,469 proteins detected, 710 were palmitoylated, accounting for 15.8% of the identified proteome (see Data S1, Fig 6). Principal component analysis (PCA) revealed consistent differences in the palmitoylome across conditions: control samples maintained at 4°C and 37°C, and samples stimulated with glycine and KCl at 37°C (STIM 37°C) (Fig 6 D). Notably, 117 proteins were differentially palmitoylated in STIM 37°C compared to control 37°C (16.4%, Fig 6 A). In part (34 of 117, 29%), these were proteins specifically palmitoylated in response to membrane depolarization by KCl and were not altered by varying temperature conditions (4°C vs 37°C). Interestingly, from 117 differentially palmitoylated proteins in STIM samples vs controls, 83.7% exhibited depalmitoylation (ratio of STIM /CTR < 1). This indicates that the primary response to KCl-mediated membrane depolarization in synaptoneurosomes is depalmitoylation of proteins. The list includes proteins implied in synaptic plasticity including synaptic vesicle fusion and neurotransmitter release proteins (complexins, dynamin-1 VAMP3, bassoon, piccolo), active cytoskeleton remodeling proteins (cofilin, drebrin, GAP43), cytoskeleton dynamics regulators (cofilin-1, complexins, drebrin, dynamin-1, DLGAP4), and proteins involved in calcium signaling (neurocalcin and neurogranin). In addition, we identified neuronal cell adhesion molecule NrCAM and neurotrimin which are known to interact between pre- and postsynaptic sites (Data S1). Stimulation with glycine and KCl induced palmitoylation changes in proteins associated with six metabolic pathways, including the citrate cycle, SNARE-mediated vesicular transport, and the synaptic vesicle cycle (Fig 6 E). Results from GO functional enrichment analysis for the STIM 37°C vs. control 37°C comparison are also shown in Fig 6 (E). In summary, our findings demonstrate that excitatory synapses may house key enzymes of the palmitoylation machinery that respond to extracellular stimuli. Specifically, stimulation with glycine and KCl led to the protein-specific addition or removal of palmitate.

**Fig 6.**
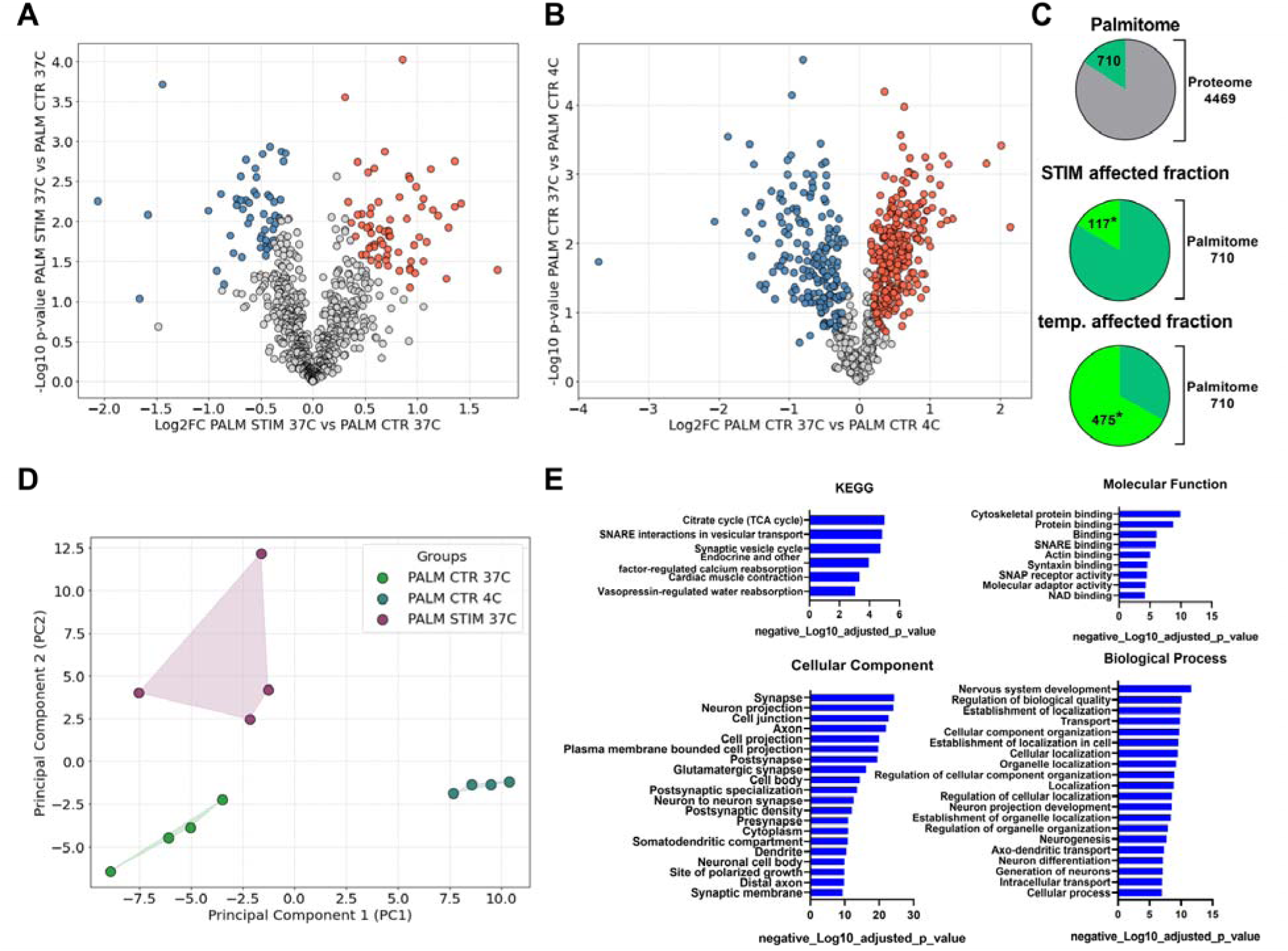
Mass spectrometry-based characterization of protein palmitoylation levels in rat hippocampal synaptoneurosomes following a 20-min stimulation (STIM) with KCl and glycine. (A) Volcano plot representing changes in protein S-palmitoylation levels in hippocampal synaptoneurosomes following stimulation with KCl (50mM) + glycine (100µM) in Mg^2+^-free solutions (PALM STIM 37°C group) relative to controls kept in the absence of these drugs in the presence of Mg^2+^ (Ctrl 37°C). A paired Student’s t-test (*n =* 4, two-sided, *S_0_* = 0.1, *FDR* = 0.05) was used to identify proteins with significant changes in palmitoylation. The X-axis shows the logl_ fold change, while the Y-axis shows the -log_10_ p-value. Red and blue dots indicate increased and decreased palmitoylation, respectively. (B) Similar Volcano plot as in (A), comparing control samples kept at 4°C versus 37°C in Mg²l_-containing aCSF. (C) Summary of numerical data presented in (A) and (B). (D) Principal component analysis (PCA) of the dataset, showing clustering of the experimental groups. (E) Results of KEGG and GO enrichment analyses for palmitoylated proteins identified in panel (A). Enriched pathways are ranked by their -log_10_ p-value. Significant GO terms (*p <* 0.05, corrected using the Benjamini-Hochberg FDR method) were identified as overrepresented.

## Discussion

In this study, we provide evidence that protein palmitoylation regulates functional short- and long-term synaptic plasticity as well as neuronal spiking of neural networks following enhanced neuron activity. We also show that following induction of synaptic plasticity, palmitoylation of synaptic proteins undergoes dynamic regulation in protein-specific and time-dependent manner. We provide the first evidence that isolated excitatory synapses contain active palmitoylation machinery responsive to external stimuli. Dynamic palmitoylation of synaptophysin and NCAM in the models used here suggests that S-PALM may be crucial for protein mobility and functional reorganization necessary for synaptic remodeling. For instance, synaptophysin depalmitoylation may enable its redistribution to perisynaptic regions, supporting vesicle recycling during LTP. Similarly, NCAM depalmitoylation may reduce synaptic adhesion, enhancing structural flexibility. The distinct roles of palmitoylation in SR versus SO synapses highlight the complexity of synaptic plasticity regulation, suggesting that different signaling pathways are engaged depending on the dendritic compartment.

### Protein-specific palmitoylation supports synaptic plasticity

It is well-established that protein palmitoylation in neurons is critically dependent on neuronal activity (4). However, global analysis of palmitoylome status of the nervous system following enhanced neuronal activity is scarce. In one recent study, stimulation of acute hippocampal slices inducing NMDAR-dependent LTP was reported to markedly increase global protein palmitoylation 30-90 min post LTP induction (37). This would suggest that palmitoylation is a unidirectional process. In contrast to this study, we do not observe a global shift in total palmitate-related signal in neuronal culture or slice homogenates following their pharmacological stimulation. In line with this, our mass spectrometry data support this finding in synaptoneurosomes (Fig. 6 A-B). One explanation for these contrasting results could be related to the different glycine concentration (100µM vs 600µM used in our study) and timing (5 min. vs 10 min. used in our study). However, in another recent mass spectrometry study, membrane fraction enriched samples of mice hippocampi were analyzed 1h post learning in a fear conditioning (aversive learning) paradigm and out of 345 palmitoylated proteins, only 35 proteins were affected (ca. 10%) (15). Similarly, the number and direction of palmitoylation for individual proteins in neurons following glutamate treatment or kainic acid-induced seizures were selective and limited to specific proteins (26). Thus, protein palmitoylation following enhanced neuronal activity appears to remain a rather subtle and target-specific process.

In this study using three different experimental models of synaptic plasticity and tissue fractions (cultures or tissue homogenates vs synaptoneurosomes) we found that presynaptic, postsynaptic or inter-synaptic proteins undergo time- and protein-dependent S-PALM. Specifically, these were synaptophysin, neurochondrin, and NCAM. With regard to presynaptic proteins, we found that unlike VAMP2 or SNAP25, synaptophysin is dynamically palmitoylated upon enhanced neuronal activity in cultured neurons, hippocampal slices and synaptoneurosomes. Synaptophysin is known for regulating neurotransmitter release and implicated in both exocytosis and endocytosis (38,39) and its expression in adult rat hippocampus seems to be similar in SR and SO (40). Interestingly, synaptophysin is involved in the clathrin-dependent endocytosis of synaptic vesicles and is required to sustain fusion events during repetitive stimulation (41). It also interacts with synaptobrevin and can regulate the fusion of the SNARE complex assembly through sequestration of the VAMP proteins (42). Additionally, synaptophysin undergoes phosphorylation upon LTP which enhances glutamate release and supports KCl-induced LTP (43). *In vivo* LTP is associated with a decrease in the number of synaptophysin labelled vesicles and increase in distance from the active zone (44). This suggests that synaptophysin requires mobility and relocation away from the active zone and toward the perisynaptic region where endocytosis takes place during LTP (44). We speculate, that depalmitoylation of synaptophysin could help increase mobility of this protein between compartments and interfering with this process adversely affects short-term presynaptic neurotransmitter release. In support of this, we also observed that protein depalmitoylation with NtBuHA resulted in reduced short-term synaptic plasticity in SR (Fig. 4). Since paired-pulse facilitation predominantly affects presynaptic release machinery (45) this supports the view that protein palmitoylation may have an important impact on presynaptic release in SR. Indeed, blocking depalmitoylating enzymes or knocking out the APT-1 gene were reported to result in increased frequency of mEPSC indicative of increased presynaptic probability of release (46). Moreover, PPT1 knockout caused a decline in the number of readily releasable synaptic vesicles (10). Thus, a picture emerges, whereby palmitoylation may support short-term synaptic plasticity in a synapse-dependent manner. On the other hand, synaptophysin contains four transmembrane domains and due to its high abundance in the presynaptic compartment it is often used as presynaptic terminal marker. Therefore over a short time period, palmitoylation could rather modify the function of synaptophysin and not its localization, yet the possible mechanism remains elusive. Importantly, a recent mass spectrometry study showed that learning in fear conditioning of mice is not associated with changes in expression and palmitoylation of synaptophysin (15). Thus, further studies are warranted to examine in detail the role of synaptophysin palmitoylation in brain plasticity, learning, and memory. Specifically, it would be insightful to investigate how the palmitoylation of synaptophysin influences vesicle recycling or neurotransmitter release.

Here, we found that palmitoylation of neurochondrin is affected by induction of neuronal plasticity in culture and slices. Neurochondrin (also known as norbin, encoded by the NCDN gene) is expressed specifically in neurons and present in somato-dendritic compartment (47). Present in the cytosol, it regulates synaptic plasticity, neuronal morphology, and signaling pathways essential for learning and memory (reviewed in (48)). Neurochondrin in dendritic spines associates with actin rather than postsynaptic density proteins such as PSD-95, which suggests a structural role in synaptic function (49). The former protein has been linked to intellectual disability and other neurodevelopmental disorders, as mutations in the NCDN gene can lead to severe cognitive and motor impairments (50). Importantly, intracellularly neurochondrin interacts with multiple proteins, of which mGluR1 and mGluR5 have been most extensively studied (48) and may function as an adaptor or scaffold protein linking membrane receptors and lipids, thus facilitating receptor trafficking and/or recycling. LTP induction results in increased neurochondrin expression which could support LTP via mGluR5. Interactor proteins of mGluRs help regulate calcium signaling, which is critical for synaptic plasticity. Neurochondrin deficiency results also in increased CaMKII activity in hippocampal lysates and impaired spatial learning (51). Importantly, in its primary sequence, neurochondrin contains 729 amino acids and multiple cysteines which could potentially undergo palmitoylation. Indeed, neurochondrin can be palmitoylated at cysteine C3 and C4, by palmitoyl-transferases ZDHHC1, ZDHHC3 and ZDHHC7 (8,52). Our study highlights the possibility that neurochondrin depalmitoylation is crucial for supporting neuronal plasticity via yet an unknown mechanism. In this regard we may speculate that such depalmitolylation prevents negative regulation of CaMKII phosphorylation by neurochondrin and promotes CaMKII activity (51). Alternatively, palmitoylation at the C-terminal region could affect an interaction of neurochondrin with proteins such as Dia1 and others (i.e. mGluR5 or CaMKII) and in this way support synaptic plasticity. The C-terminus of neurochondrin was found to interact with diaphanous related formins like Dia1 and affect structural plasticity by regulating neurite outgrowth (53).

Neural cell adhesion molecule NCAM mediates cell-cell and cell-matrix adhesion and plays a key role in neurite outgrowth, synaptogenesis, and synaptic plasticity (54). NCAM contains 3 cysteines that undergo palmitoylation. This modification helps NCAM target lipid rafts, facilitates NCAM-mediated signaling and neurite outgrowth (55). It was shown previously that NCAM140 and NCAM180 are covalently modified by thioester-linked palmitate following fibroblast growth factor 2 (FGF2) - stimulated neurite outgrowth (56) and NCAM was identified as substrate for ZDHHC3 and ZDHHC7 (8). Palmitoylation of NCAM may also help in its endocytosis and recycling to the plasma membrane (57). A presence of NCAM blocking antibody or NCAM knockout both have negative effects on the early phase of synaptic potentiation lasting minutes (58). In this study NCAM palmitoylation was decreased 20 minutes after induction of synaptic plasticity with cLTP in neuronal cultures or stimulation of synaptoneurosomes. The role of rapid NCAM depalmitoylation in supporting synaptic plasticity remains unclear. We propose that NCAM depalmitoylation increases its mobility between compartments and reduces the proportion of NCAM anchored in the membrane toward the extracellular space. This reduction in NCAM-mediated synaptic adhesion and interactions with extracellular matrix components, including heparan proteoglycans, is thought to enhance the structural flexibility required for synaptic plasticity.

### Synaptoneurosomes and protein depalmitoylation upon LTP

One of the most surprising findings of this study is observation that exposing live, extracted excitatory synapses to a temperature change or glycine and KCl (STIM) resulted in protein-dependent dynamic palmitoylation occurring within minutes. The results are highly reproducible across all samples analyzed (Fig. 6C). In this study we used previously established protocols that allowed us to prepare live synaptoneurosomes capable of undergoing NMDAR-dependent LTP. In support of this view, we showed increased expression of pERK, a marker of neuronal plasticity, following glycine and KCl stimulation using Western blotting (24). In addition, with mass spectrometry we detected increased expression of 90 other proteins post STIM in agreement with the idea of local translation of proteins in the synapse (35). An overview of the results indicate the presence of proteins involved in pathways such as synaptic vesicle cycling, neurotransmitter release, and cellular metabolism (Data S1). The identified proteins also contribute to cellular structural organization, ion transport, and nervous system development. Thus, this model provides a viable way to study the molecular machinery of the isolated excitatory synapses.

Previous studies implied that in neurons several enzymes including ZDHHC1, ZDHHC2, ZDHHC5 and ZDHHC8 may be present in dendrites or synapses (8), while ZDHHC3, ZDHHC7 and ZDHHC12 were found in somatic Golgi. ZDHHC17 was additionally associated with several vesicular structures, including the Golgi apparatus as well as sorting/recycling and late endosomal structures and was found in presynaptic terminal in drosophila (59). Interestingly, overexpression of some ZDHHCs led to their discovery in Golgi outposts (60). In this study using mass spectrometry of synaptoneurosomes we found the presence of only two ZDHHCs (ZDHHC5 and ZDHHC17) and all known members of depalmitoylating machinery including APT1/2, PPT1, and ABHDs (see also (26)). We did not detect Golgi marker proteins in synaptoneurosomal fractions (i.e. GM130, SialT2, TGN38 (Data S1) (61)). The use of live synaptoneurosomes excludes the possibility of extensive ultracentrifugation to obtain ultrapure synaptic vesicles. However, our protocol for generating live synaptoneurosomes results in a significant enrichment of both pre- and postsynaptic markers and a depletion of cytosolic and nuclear markers in the SN fraction as compared to the homogenate (62). Therefore palmitoylation in synaptoneurosomes could be mediated primarily by non-Golgi located ZDHHCs like ZDHHC5 while depalmitoylation enzymes seem to be overrepresented. Interestingly, mass spectrometry comparison of the palmitoylomes of samples kept at 4°C vs 37°C for 30 minutes showed a distinct pattern of palmitoylation of hundreds of proteins (Fig. 6). Palmitoylation as an enzymatic process is expected to strongly depend on the temperature and thioester bond is stable and covalent. Thus, one explanation for this results is the difference in enzymatic activity in the preparations further supporting a view that palmitoylation machinery is present in the excitatory synapses and responsive to external stimuli.

It was previously shown that PPT1 is present in the synaptic compartment of the rat neurons and high levels of PPT1 enzyme activity was found in the synaptic cytosol (63). In addition, approximately 10% of palmitoylated synaptic proteins were reported as a substrate of PPT1. In this study, among palmitoylated proteins we found that neural cell adhesion molecule (Nrcam), neurotrimin and sodium/potassium-transporting ATPase subunit beta-2 were significantly depalmitoylated 20 min post glycine + KCl treatment. These proteins were reported substrates to PPT1 (63). Thus, one explanation of our results is that PPT1 activity in excitatory synapses may be triggered by external stimulation.

While ABE on synaptoneurosomes showed massive depalmitoylation of proteins such as synaptophysin, NCAM, and PSD95 (Fig. 5), quantification by mass spectrometry confirmed only PSD95 (Table 1). This discrepancy may arise due to significant differences between the techniques used. First, shotgun proteomics identifies tryptic peptides, whereas Western blot relies on antibody specificity, which can vary. Additionally, unlike Western blotting, mass spectrometry involves multiple normalization steps. We subtracted protein intensities in PALM samples not treated with hydroxylamine from the corresponding hydroxylamine-treated samples (to control for nonspecific binding), then subtracted protein intensities in input samples from PALM samples (to correct for potential protein level differences), and subtracted the median protein intensity within PALM samples to normalize acyl-proteome levels. Only then did we calculate protein level differences between PALM sample groups. Furthermore, additional post-translational modifications (PTMs) can affect protein quantification by both techniques. PTM-modified peptides may not appear in proteomics unless specifically queried, meaning that if a protein is quantified by a small number of peptides, PTM-level variations will influence protein quantification. Therefore, the results of mass spectrometry are most informative when analyzed from a population perspective, rather than focusing on individual proteins.

### Calcium signaling and the dynamic regulation of protein palmitoylation in neuronal plasticity

It is well established that in neurons, protein palmitoylation heavily depends on neuronal activity, and intracellular cascades mediating the transduction of extracellular stimuli to palmitoylation machinery activity are beginning to be unraveled (4). However, a key question arises: how do external stimuli translate into the control of protein palmitoylation on demand in neurons? In this study, we induced NMDAR-dependent forms of synaptic plasticity to mimic those occurring *in vivo* (16,18). NMDARs regulate Ca² entry into the postsynaptic compartment. Therefore, in a simplified view, palmitoylation may be regulated by pathways that are heavily dependent on Ca². Indeed, palmitate cycling on PSD-95 was shown to be regulated by Ca² influx through NMDA receptors and to be absent in Ca² -free solutions (64). Additionally, NCAM was shown to interact with calmodulin, a Ca² -binding protein. Translocation of NCAM to the ER and from the ER membrane to the cytoplasm, and its import from the cytoplasm to the nucleus are calmodulin- and calcium-dependent processes (65). Moreover, external stimuli such as fibroblast growth factor 2 (FGF-2), which is known to promote LTP in the hippocampus, have been shown to induce NCAM palmitoylation (56). Interestingly, FGF-2 regulates NMDAR-mediated Ca² entry (66). Thus, dynamic changes in intracellular Ca² seem to directly translate to protein palmitoylation. However, it remains unknown how cells manage to regulate each protein individually, resulting in the complex, protein-specific patterns of palmitoylation observed in this study (e.g., neurochondrin vs. PSD-95). It is crucial to further explore the molecular signals regulating protein palmitoylation in neuronal plasticity in greater detail.

Interestingly, the direction of palmitoylation for individual proteins varied depending on the experimental model, cellular fraction, and timing after LTP induction. Results were consistent across slices and synaptoneurosomes but showed mixed outcomes in neuronal cultures (Fig. 5G). A possible explanation is that the chemical LTP protocol used in cultures broadly activates plasticity mechanisms, including enhanced neuronal activity, adenylyl cyclase stimulation, and increased cAMP and cGMP levels via phosphodiesterase inhibition. In contrast, synaptic NMDAR-dependent LTP was induced in slices and synaptoneurosomes where synapses were stimulated with glycine in the absence of magnesium ions. This suggests that the engagement of palmitoyltransferases and thioesterases may differ depending on the molecular pathways activated to induce plasticity. Future studies could investigate these differences further by directly comparing the models used in this study. Notably, in slice homogenates, most proteins showed increased palmitoylation following LTP induction, whereas synaptoneurosomes exhibited preferential depalmitoylation. We hypothesize that fractionation of the nervous tissue to the level of synaptoneurosomes alters the balance of palmitoyltransferases and thioesterases, shifting the equilibrium of S-palmitoylation. Supporting this, mass spectrometry revealed a significantly higher abundance of depalmitoylating enzymes in synaptoneurosomes, suggesting that depalmitoylation would dominate under these conditions. Future studies could determine the activity levels of ZDHHCs and depalmitoylases during LTP. Currently the tools to determine that are lacking.

### Palmitoylation supports short-and long-term synaptic plasticity in a synapse-specific manner

In this study, we report that the deacylation agent NtBuHA disrupted long-term synaptic plasticity in excitatory synapses of apical dendrites of CA1 pyramidal neurons (Fig. 4). This observation aligns with a recent report showing that inhibition of hippocampal palmitoyl acyltransferase activity with 2-BP impairs *in vivo* hfsLTP in the CA1 region of the hippocampus (14). Additionally, transgenic animals lacking ZDHHC2, ZDHHC17, or PPT1 were reported to have impaired LTP (11,67,68). Thus, palmitoylation appears to be indispensable for supporting synaptic plasticity. However, we also show that in the SO, NtBuHA did not affect the time course of hfsLTP nor the short-term synaptic plasticity (Fig. 4). The underlying mechanisms remain elusive. Nevertheless, we previously demonstrated that while excitatory synapses in the SO and SR require NMDAR activity as a critical component for LTP induction (32), they differ significantly in the downstream molecular machinery supporting synaptic potentiation. Specifically, protein kinase C, RhoGTPases, and matrix metalloproteases were crucial for synaptic potentiation in SR, whereas SO synapses required Src kinases and dopamine receptors (32). Furthermore, SO is devoid of TrkB, the receptor for BDNF (69). It has also been reported that SO synapses rely on L-type voltage-gated calcium channels for dopamine-induced LTP, in contrast to SR synapses, which depend on BDNF signaling (70). Thus, S-palmitoylation may regulate only certain pathways required for NMDAR-dependent plasticity. Pyramidal neurons, with their basal and apical dendrites, play a crucial role in information processing. However, it remains unclear why excitatory synapses in the dendritic trees express different molecular mechanisms of plasticity. One possible explanation is that the spatial segregation of inputs and the implementation of different forms of synaptic plasticity in basal versus apical dendrites enable pyramidal neurons to enhance their computational capacity, which is vital for information processing. Future studies could benefit from investigating the differences between these two types of neighboring excitatory synapses to further elucidate this issue. For example, it would be valuable to explore why SR and SO synapses have distinct palmitoylation dependencies, using molecular inhibitors that target specific pathways or by silencing genes involved in palmitoylation machinery (i.e. ZDHHCs). Additionally, investigating S-palmitoylation’s role in other hippocampal connections, such as mossy fiber to CA3 pyramidal neurons, known for expressing NMDAR-independent forms of LTP, could be illuminating (33).

### Palmitoylation in network plasticity and memory formation

This study provides evidence that protein palmitoylation is crucial for the temporal reorganization of neuronal spiking in neural networks following episodes of enhanced neuronal activity (Fig. 1). These network changes may be explained by the negative effects of protein depalmitoylation on synaptic plasticity *in vitro*. Thus, protein palmitoylation not only supports local synaptic plasticity but also plays a critical role in the temporal organization of neuronal spiking in networks, ultimately contributing to engram formation. Memory formation occurs when highly excitable neurons display coordinated activity during encoding, and in some cases, it does not require synaptic plasticity (reviewed in (71)). Supporting the role of palmitoylation in learning and memory, a recent study showed that infusion of 2-BP into the hippocampus disrupted the acquisition and maintenance of spatial memories, but not the recall of these memories (14). Similarly, transgenic animals lacking ZDHHC2, ZDHHC9, or PPT1 exhibited impaired memory formation (11,67,68). Conversely, learning-induced changes in the brain’s palmitoylome further support its involvement in memory processes (15,46). Since the depalmitoylating agent NtBuHA did not affect the basal temporal organization of spikes or spiking frequency *in vitro* before electrical stimulation, we did not further study depalmitoylation’s impact on neural intrinsic excitability. However, we cannot exclude the possibility that palmitoylation also affects other nonsynaptic forms of neural plasticity. For instance, enhanced neuronal activity could lead to permanent changes in neuronal excitability, associated with altered passive membrane properties and ion channel expression, in the absence of synaptic gain (72).

In this study, we employed an electrical stimulation protocol previously demonstrated to induce network plasticity (73). This paradigm was selected for two primary reasons. First, it is an elegant approach, as neurons remain in the incubator and are recorded without direct intervention by the experimenter. Second, our preliminary attempts to induce network plasticity using chemically induced long-term potentiation (cLTP) proved unreliable, yielding inconsistent results of either potentiation or depression of neuronal spiking (data not shown). Additionally, the cLTP protocol required removing the microelectrode array (MEA) from the incubator, exchanging the medium, and mixing, which could influence neuronal spiking within a short time frame.

### Effects of palmitoylation inhibitors on basal transmission

In our experiments, 1 mM NtBuHA did not negatively affect basal fEPSPs in SR and SO (Fig. 4) or neuronal spiking in cultures on MEA (Fig. 1). This aligns with studies showing that NtBuHA (at 100 μM) did not affect the palmitoylation levels of PSD-95, AMPAR, or NMDAR subunits in cultured neurons (46). However, conflicting reports exist. For example, acute application of NtBuHA to hippocampal slices was reported to downregulate excitatory synaptic transmission in CA1 pyramidal neurons by reducing AMPAR surface expression (28). Reduced palmitoylation of PSD-95 and AKAP79/150 was implicated in that process (28). While another inhibitor, 2-BP, did not affect mEPSC frequency or amplitude in hippocampal slices (46), it altered basal excitatory and inhibitory synaptic transmission in basolateral amygdala neurons (74). Furthermore, *in vivo* application of high doses of 2-BP reportedly downregulated basal fEPSPs in the CA1 region of the hippocampus (14). Therefore, whether S-PALM inhibitors affect basal synaptic transmission remains a matter of debate.

## Materials and methods

### Primary neuronal cultures

All procedures on animals were carried out following the guidelines established and approved by the Polish Ethical Committee on Animal Research. Dissociated hippocampal cultures were prepared from postnatal day 0 (P0) Wistar rats as described previously (75). For click chemistry experiments, cells were plated on 13-mm-diameter coverslip (Karl Hecht glassware factory, Germany) coated with 50 µg/ml poly-D-lysine (Merck, Poland) and 2.5µg/ml laminin (Roche, Poland ) at a density of 80,000 cells per coverslip. For ABE method, neurons were cultured in 6-well plates coated with 50 µg/ml poly-D-lysine at a density of 450,000 cells. The cultures were kept at 37 °C in 5% CO2 in a humidified incubator. The experiments were performed at 14-15 day *in vitro* (DIV).

### Cytotoxicity test

For verification of cytotoxicity of drugs used in this study, we used the CytoTox 96® Non-Radioactive Cytotoxicity Assay (cat nr G1780, Promega Corporation, USA) and followed producer guidelines. The test relied on quantitative measurement of lactate dehydrogenase (LDH) released from cells with enzymatic assay, which results in the conversion of a tetrazolium salt (iodonitro-tetrazolium violet; INT) into a red formazan product. The intensity of color formed was proportional to the number of lysed cells. Visible wavelength absorbance data were collected at 490nm using a standard 96-well plate reader. 50µL of hippocampal cultures media (3 independent cultures, experimental LDF release) were compared with medium obtained from wells where cells were subjected to complete lysis (maximum LDH release) or fresh medium (negative control).

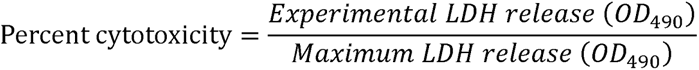

### Synaptoneurosomes

Synaptoneurosomes have been prepared as we described previously (76) with modifications. Briefly, artificial cerebrospinal fluid (aCSF) (containing in mM: 87 NaCl, 2.5 KCl, 1.25 NaH_2_PO_4_, 25 NaHCO_3_, 0.5 CaCl_2_, 10 MgSO_4_, 20 glucose, 75 sucrose) was aerated with an aquarium pump for 30 min at 4°C. Next, the pH was adjusted to 7.4 with dry ice. The buffer was supplemented with 1 × protease inhibitor cocktail cOmplete EDTA-free (Roche; 05056489001). Next, one month old Wistar rats were euthanized with cervical dislocation and hippocampi were dissected. The hippocampi were cut in halves along the transverse section then each part was homogenized in 1.5 ml of aCSF using Dounce homogenizer with 10–12 strokes. All steps were kept ice-cold to prevent stimulation of synaptoneurosomes. For each sample, homogenates from hippocampi of 2 rats were diluted in an ice-cold aCSF buffer to the final volume of 18 ml. The samples were loaded into 20 ml syringe and passed through a series of aCSF-pre-soaked nylon mesh filters with a pore-size of consecutively 100, 60, 30 and 10 µm (Nylon Net Filters; Merck Millipore) to 50 ml polypropylene tube in a cold room. The filtrate was centrifuged at 1000 × g for 15 min at 4⍰C (Eppendorf S-4-72 swinging bucket rotor). The pellet was resuspended in 3 ml of ice-cold aCSF buffer with protease inhibitors and the sample was divided into 3 parts. Synaptoneurosomes were stimulated as described previously (36) with modifications resulting in three sample types: unstimulated 4⍰C and 37⍰C control samples and stimulated 37⍰C. Briefly, the suspensions of synaptoneurosomes were centrifuged at 1000 × g for 15 min at 4⍰C. The 4⍰C and 37⍰C control samples were resuspended in the aCSF solution (containing in mM: 125 NaCl, 2.5 KCl, 1.25 NaH_2_PO_4_, 25 NaHCO_3_, 2.5 CaCl_2_, 1.5 MgSO_4_, 20 glucose). The 4⍰C control sample was left on ice for for 30 min and then frozen. The 37⍰C control sample was supplemented with strychnine (25µM) and incubated for 30 min in 37⍰C with shaking and then frozen. Simultaneously, the stimulated sample was resuspended in magnesium-free aCSF solution (containing in mM: 125 NaCl, 26 NaHCO_3_, 1.6 NaH_2_PO_4_, 2.5 CaCl_2_, 5 KCl and 10 glucose), supplemented with strychnine (25µM) and glycine (100µM) and incubated for 10 min in 37⍰C with shaking followed by application of high KCl (50 mM final concentration) for another 20 minutes and then frozen.

### Multi electrode arrays

Multielectrode arrays containing 60 micro-electrodes of 30 µm diameter and 200 µm spacing (60PedotMEA200/30iR-Au, Multichannel Systems, Germany) were autoclaved, covered with fetal bovine serum and left for 30 minutes to become hydrophilic. Following wash with sterile water, plates were coated with PDL (0.05mg/ml) overnight at 37°C and then coated again with laminin (20µg/ml) and left overnight at 37°C. Neurons were plated at density 1 million cells / plate in Minimal Essential Medium (MEM, Gibco, Thermofisher, Poland) for 2h and subsequently cultured in Neurobasal Medium (Gibco, Thermofisher, Poland) with fluorinated ethylene-propylene membrane covers (ALA MEA sheets, ALA Scientific Instruments Inc., USA) for up to 21 days. All recordings of spontaneous neuronal network activity were made inside the incubator at 37°C with MEA2100-Mini System (Multichannel Systems, Germany). Recording electrodes were considered active if spiking frequency was > 0.1 Hz. Only plates with > 60 % of active electrodes were considered for further experiments. Signals were band-pass filtered at 300-5000 Hz and digitized at 25kHz. All recordings were made in 5 minute-long epochs every 15-30 minutes. Stimulation of the electrodes on the MEA was performed with AT-IN protocol as described previously (77). Briefly, following basal activity recordings, stimulation was applied on a pair of two active electrodes recording neuronal spiking in two distant areas of the plate. A low-frequency stimulation consisting of 120 stimuli at 1Hz was applied to one single electrode and theta burst stimulation (5 stimuli applied at 100Hz, every 50ms) to another electrode. Single pulse was applied in phase with the burst and such associative stimulation was applied for 2 minutes. Each pulse was bi-phasic, ±800 mV (positive phase first) and lasted for 500 µs. Spontaneous neuronal activity was recorded for 60 minutes post stimulation. Spike detection was set at 6 times standard deviation of noise. Data analysis was performed in software provided by the manufacturer and custom program written in Python.

### Acute brain slices and electrophysiology

Acute hippocampal brain slices from 1.5-2 month old rats were prepared according to the protocol described previously. Briefly, hippocampi from one hemisphere were dissected and cut into 350 µm thick slices using a vibratome (5100mz, Campden Instruments, USA) in oxygenated (95% O2, 5% CO2) ice-cold buffer with pH=7.4 containing in mM 92 NMDG, 2. KCl, 1.2 NaH_2_PO_4_, 30 NaHCO_3_, 0.5 CaCl_2_, 10 MgSO_4_*7H_2_O, 20 HEPES, 25 glucose, 2 thiourea, 5 sodium ascorbate, 3 sodium pyruvate. Slices were recovered in solution containing in mM: 92 NaCl, 2.5 KCl, 1.2 NaH_2_PO_4_, 30 NaHCO_3_, 2 CaCl_2_, 2 MgSO_4_*7H_2_O, 20 HEPES, 25 glucose, 2 thiourea, 5 sodium ascorbate, 3 sodium pyruvate for 15 min (32 °C). Finally, slices were stored in oxygenated (95% O2, 5% CO2) artificial cerebrospinal fluid (aCSF) that contained in mM: 124 NaCl, 2.5 KCl, 1.2 NaH_2_PO_4_, 24 NaHCO_3_, 2 CaCl_2_, 2 MgSO_4_*7H_2_O, 5 HEPES, 12.5 glucose, pH 7.4. Recordings were made in aCSF after 2 h of slice recovery. Test stimuli (0.3 ms, 0.1 Hz) were delivered using a stimulator (A385, World Precision Instruments, USA) and a concentric bipolar electrode (FHC, Bowdoin, ME USA) placed either in SR or SO. fEPSPs were recorded with glass micropipettes that were filled with aCS*F (*1-3 M Ω resistance). Input–output (I–O) relationships were built for fEPSPs amplitudes upon monotonically increasing the stimuli in the range of 0–300 µA (13 points, applied once at 0.1 Hz). Baseline stimulation was set at 0.1 Hz, and for baseline and paired-pulse stimulation protocols (interstimulus interval 50 ms), the stimulation strength was set to 40% of the maximum fEPSP amplitude. Signals were acquired with Sutter Double IPA amplifier (Sutter Instruments, USA)

### Acyl Biotin Exchange method

To analyze changes in protein palmitoylation levels acyl-biotin exchange assay (ABE) was used. To extract protein from rat hippocampal tissue slices, synaptoneurosomes or primary hippocampal neurons at 14 DIV the material was lysed and homogenized using Dounce homogenizer in a buffer containing in mM: 50 Tris HCl (pH 7.5), 150 NaCl, 1 EDTA and 4% SDS, 1% Triton X-100. Proteins were reduced with 10 mM TCEP (tris(2-carboxyethyl)phosphine) for 30 min at room temperature (RT) and subsequently, the samples were incubated for 16h at 4°C with 50mM N-ethylmaleimide (NEM) to block free thiol groups. Next, proteins were precipitated using chloroform-methanol extraction and subsequently washed three times with 96% ethanol to remove unreacted NEM. The pellets were resuspended in the same buffer as used previously. The fraction of each sample was moved to a separate tube to create pooled-within-condition negative control samples. The remaining volume underwent the acyl-biotin exchange reaction (‘positive samples’). Both positive and negative control samples were treated with 400 µM thiol-reactive biotinylation reagent HPDP-biotin (N-[6-(biotinamido)hexyl]-30-(20-pyridyldithio)propionamide) for 1.5h at RT. Simultaneously, only the positive samples were treated with 1M hydroxylamine to cleave thioester-linked palmitoyl moieties and expose newly formed-thiols to HPDP-biotin. Subsequently, proteins were precipitated with chloroform-methanol method, washed three times with ethanol, and re-dissolved in a buffer with a lower detergent content (containing in mM: 50 Tris HCl (pH 7.5), 150 NaCl, 1 EDTA and 0.5% SDS, 0.2% Triton X-100). At this stage the samples were subjected to SDS-PAGE and Western blotting to analyze the global state of S-palmitoylation (Stain-free total protein visualization was used as a loading control) or processed further in order to isolate enriched S-palmitoylated protein fraction. To this end, equal amounts of protein were taken from each sample and incubated with Pierce™ High Capacity NeutrAvidin™ Agarose beads for 2h at RT. Subsequently, the beads were washed 6 times with a buffer containing in mM: 50mM Tris (pH7.7), 600 NaCl, EDTA). To elute the enriched S-palmitoylated proteins fraction the beads were incubated with 50 mM Tris buffer (pH 7.7) with 1% β-mercaptoethanol for 1.5 h at 37°C. Finally, aliquots of the samples were subjected to SDS-PAGE and Western blotting to visualize and analyze palmitoylation level changes to individual proteins.

### Western blot analysis

Protein concentration in the samples was determined using a Pierce™ BCA protein Assay kit (Thermo Scientific). The samples of equal protein content or equal volume (in case of isolated enriched S-palmitoylated protein fraction) were supplemented with with a 6x Lammli sample buffer and heated at 95 °C for 5 min. Subsequently, the samples were subjected to a standard SDS-PAGE in TGX Stain-Free FastCast Acrylamide 10% gels (Bio-Rad) and then transferred to PVDF membranes (Immobilon-FL, Millipore) using an electrophoretic transfer system (Trans-Blot Turbo, Bio-Rad). The non-specific binding sites were blocked with either 10% skimmed milk or EveryBlot Blocking Buffer (Bio-Rad) for 1 h at RT. Then, the blots were incubated overnight at 4°C with the primary antibodies (Table below). After incubation, the blots were washed 5x with TBS-T and subsequently incubated with secondary antibodies (Table) for 1.5 hours at RT. Blots were visualized using ECL substrate (Clarity and Clarity Max, Bio-rad) and scanned using a Chemidoc image analyzer (Bio-Rad). The molecular weights of immunoreactive bands were estimated on the basis of the migration of molecular weight markers (Thermo Fisher Scientific). Signal intensity analysis was conducted using ImageLab software (Bio-Rad).

| Antibodies |  |  |  |  |  |  |
| --- | --- | --- | --- | --- | --- | --- |
| Antibody | Catalogue Number | Host | Blocking Agent | Primary Dilution | Manufacturer | Secondary Dilution |
| Synaptophysin (37 kDa) | MAB329 | Mouse | EveryBlot | 1:2500 | Merck | Anti-mouse 1:5000 |
| SNAP-25 (25 kDa) | 111004 | Guinea Pig | EveryBlot | 1:2500 | SYSY antibodies | Anti-guinea pig 1:3000 |
| Phospho-p44/42 MAPK (Erk1/2) (44 kDa) | 4377S | Rabbit | Milk | 1:1000 | Cell Signalling | Anti-rabbit 1:5000 |
| PSD-95 (100 kDa) | ab18258 | Rabbit | Milk | 1:2500 | Abcam | Anti-rabbit 1:5000 |
| Arc (Arc/Arg3.1) (50 kDa) | 156003 | Rabbit | Milk | 1:1000 | SYSY antibodies | Anti-rabbit 1:5000 |
| NCAM (120-220 kDa) | A309700 | Rabbit | Milk | 1:1000 | Antibodies.com | Anti-rabbit 1:5000 |
| Neurochondrin (75 kDa) | A53729 | Rabbit | Milk | 1:500 | Antibodies.com | Anti-rabbit 1:5000 |
| VAMP2 (15kDa) | ab3347 | Rabbit | EveryBlot | 1:1000 | Abcam | Anti-rabbit 1:5000 |
| GluR1 (100 kDa) | ABN241 | Rabbit | EveryBlot | 1:1000 | Merck | Anti-rabbit 1:5000 |

### Click chemistry

Neuronal cell cultures were grown on coverslips and on 13 DIV received alkyne-palmitic acid (50 μM) dissolved in dimethyl sulfoxide (Sigma Aldrich). Additionally, some cells received NtBuHA dissolved in water (1 mM) or solvent and were incubated overnight. On 14 DIV cells were washed three times with phosphate-buffered saline (PBS), fixed in cold methanol for 10 min at -20°C, and permeabilized in 0.1% Triton X-100 in PBS for 5 min. The click-chemistry reaction involved 1 h incubation at room temperature with a mixture of Oregon Green 488 azide (0.1 mM) (Thermo Fisher Scientific), TCEP (1 mM), and CuSO4 (0.1 mM). Next, the cells were washed five times with PBS, and the coverslips were mounted in Fluoromount G anti-quenching medium. Images of neurons were acquired using a Zeiss LSM780 laser scanning confocal microscope with a Plan Apochromat 63×/1.4 oil immersion objective using a 488 nm diode-pumped solid-state laser at a pixel count 1024 × 1024 at 16 bit depth. A series of z-stacks were collected for each stimulation with a 1 μm step size. The fluorescence intensity was determined using Fiji open source image processing package (78).

### Proteomic sample preparation, LC-MS/MS measurements, and data analysis

INPUT SAMPLES were subjected to chloroform/methanol precipitation, and the resulting protein pellets were washed twice with methanol. The protein pellets were air-dried for ∼10 min and resuspended in 100 mM HEPES (pH 8) by vigorous vortexing and sonication. Proteins were then reduced and alkylated with 10 mM tris(2-carboxyethyl)phosphine hydrochloride (TCEP)/40 mM chloroacetamide (CAA) and digested with trypsin in a 1:30 (w/w) enzyme-to-protein ratio at 37°C overnight. Digestion was terminated by the addition of trifluoroacetic acid (TFA) to a 1% final concentration. The resulting peptides were labelled using an on-column TMT labelling protocol (79). TMT-labelled samples were compiled into a single TMT sample and concentrated using a SpeedVac concentrator. Peptides in the compiled sample were separated into fractions with basic reversed-phase using the Pierce High pH Reversed-Phase Peptide Fractionation Kit (Thermo Fisher Scientific). PALM SAMPLES were dried using a SpeedVac concentrator, then resuspended in 100 mM HEPES (pH 8) by vigorous vortexing and sonication. Proteins were reduced and alkylated with 10 mM TCEP/40 mM CAA and digested with trypsin in a 1:30 (w/w) enzyme-to-protein ratio at 37°C overnight. Digestion was terminated by the addition of trifluoroacetic acid (TFA) to a 1% final concentration. The resulting peptides were labelled using an on-column TMT labeling protocol. TMT-labelled samples were compiled into a single TMT sample and concentrated using a SpeedVac concentrator. Prior to the liquid chromatography (LC)-MS measurement, the peptide fractions were resuspended in 0.1% TFA and 2% acetonitrile in water. Chromatographic separation was performed on an Easy-Spray Acclaim PepMap column (50 cm length × 75 µm inner diameter; Thermo Fisher Scientific) at 55°C by applying 120 min acetonitrile gradients in 0.1% aqueous formic acid at a flow rate of 300 nl/min. An UltiMate 3000 nano-LC system was coupled to a Q Exactive HF-X mass spectrometer via an easy-spray source (all Thermo Fisher Scientific). The Q Exactive HF-X was operated in TMT mode with survey scans acquired at a resolution of 60,000 at m/z 200. Up to 15 of the most abundant isotope patterns with charges 2-5 from the survey scan were selected with an isolation window of 0.7 m/z and fragmented by higher-energy collision dissociation with normalized collision energies of 32, while the dynamic exclusion was set to 35 s. The maximum ion injection times for the survey scan and dual MS (MS/MS) scans (acquired with a resolution of 45,000 at m/z 200) were 50 and 120 ms, respectively. The ion target value for MS was set to 3e6 and for MS/MS was set to 1e5, and the minimum AGC target was set to 1e3. Prior to the liquid chromatography (LC)-MS measurement, the peptide fractions were resuspended in 0.1% TFA and 2% acetonitrile in water. Chromatographic separation was performed on an Easy-Spray Acclaim PepMap column (50 cm length × 75 µm inner diameter; Thermo Fisher Scientific) at 55°C by applying 120 min acetonitrile gradients in 0.1% aqueous formic acid at a flow rate of 300 nl/min. An UltiMate 3000 nano-LC system was coupled to a Q Exactive HF-X mass spectrometer via an easy-spray source (all Thermo Fisher Scientific). The Q Exactive HF-X was operated in TMT mode with survey scans acquired at a resolution of 60,000 at m/z 200. Up to 15 of the most abundant isotope patterns with charges 2-5 from the survey scan were selected with an isolation window of 0.7 m/z and fragmented by higher-energy collision dissociation with normalized collision energies of 32, while the dynamic exclusion was set to 35 s. The maximum ion injection times for the survey scan and dual MS (MS/MS) scans (acquired with a resolution of 45,000 at m/z 200) were 50 and 120 ms, respectively. The ion target value for MS was set to 3e6 and for MS/MS was set to 1e5, and the minimum AGC target was set to 1e3. The data were processed with MaxQuant v. 1.6.17.0 and the peptides were identified from the MS/MS spectra searched against Uniprot Rat Reference Proteome (UP000002494) using the built-in Andromeda search engine. Raw data related to PALM SAMPLES and INPUT SAMPLES were processed together using MaxQuant software. Reporter ion MS2-based quantification was applied with reporter mass tolerance = 0.003 Da and min. reporter PIF = 0.75. Cysteine carbamidomethylation was set as a fixed modification and methionine oxidation, glutamine/asparagine deamidation as well as protein N-terminal acetylation were set as variable modifications. For in silico digests of the reference proteome, cleavages of arginine or lysine followed by any amino acid were allowed (trypsin/P), and up to two missed cleavages were allowed. The FDR was set to 0.01 for peptides, proteins and sites. Match between runs was enabled. Other parameters were used as pre-set in the software. Reporter intensity corrected values for protein groups were loaded into Perseus v. 1.6.10. Standard filtering steps were applied to clean up the dataset: reverse (matched to decoy database), only identified by site, and potential contaminant (from a list of commonly occurring contaminants included in MaxQuant) protein groups were removed. Reporter intensity corrected values were log2 transformed and protein groups with values for all PALM samples were kept. The values recorded for the PALM samples not treated with hydroxylamine were subtracted from the values recorded for the PALM samples subjected to hydroxylamine treatment. Values recorded for the INPUT samples were normalized by median subtraction within TMT channels. The values recorded for the INPUT samples were subtracted from the values recorded for the corresponding PALM samples. Median was then subtracted within TMT channels. To determine which proteins were differentially palmitoylated following temperature shift/stimulation. Student’s t-tests (2-sided, permutation-based FDR□=□0.05, S0□=□0.1, n = 4) were performed on the groups of PALM samples. In addition, Student’s t-tests (2-sided, permutation-based FDR□=□0.05, S0□=□0.1, n = 4) were performed on the groups of INPUT samples to determine possible changes in protein levels following temperature shift/stimulation. For lists of proteins identified and quantified in these experiments, see Data S1. This dataset was deposited to the ProteomeXchange Consortium via the PRIDE partner repository with the dataset identifier PXD058417. To characterize palmitoylated proteins in rat hippocampal synaptoneurosomes we used TMT-based quantitative proteomics. KEGG and GO functional protein enrichment analyses were performed using the web-based tool ShinyGO 0.81. The 116 differentially palmitoylated proteins identified in synaptoneurosomes following KCl stimulation (mapped to 101 unique rat genes, or 87%) were queried against the Rattus norvegicus hippocampal proteome database, specifically proteins identified with TMT labeling (6,359 proteins (80)). Pathway enrichment analysis (KEGG) and enrichment for GO molecular function, cellular component, and biological process terms were limited to top 20 results. Enriched pathways were ordered by the negative logarithm of their p-value. All significant GO terms with *p <* 0.05 (corrected for multiple testing using the Benjamini-Hochberg false discovery rate method) were selected as overrepresented.

## Acknowledgments

The authors would like to thank colleagues from the Nencki Institute of Experimental Biology (Warsaw, Poland) Wing-Sze Tse, Krystian Bijata (Laboratory of Cell Biophysics) and Mark Hunt (Laboratory of Neuroinformatics) for consultations. We thank Łukasz Chrobok (Department of Neurophysiology and Chronobiology, Institute of Zoology and Biomedical Research, Faculty of Biology, Jagiellonian University, Krakow, Poland) and Wojciech Pokrzywa (Laboratory of Protein Metabolism, International Institute of Molecular and Cell Biology in Warsaw) for discussions. We thank Kasia Radwańska (Laboratory of Molecular Basis of Behavior, Nencki Institute of Experimental Biology, Warsaw, Poland) and Łukasz Szewczyk and Marta Wiśniewska (Laboratory of Molecular Neurobiology, Centre of New Technologies, University of Warsaw, Poland) for providing anti-Arc and anti-VAMP2 antibodies, respectively. We thank the Laboratory of Imaging Tissue Structure and Function at Nencki Institute of Experimental Biology (Warsaw, Poland) for providing equipment for confocal microscopy imaging.

## Funding

National Science Center grant SONATA BIS 2019/34/E/NZ4/00387 (TW)

## Author contributions

Conceptualization: AP, TW

Data curation: AP, TW

Formal analysis: AP, RI, RD, EK, IF, RS, TW

Funding acquisition: TW

Investigation: AP, RI, AB, JM, IF, PW, NM, AW, MR, TW

Methodology: AP, RI, AB, JM, IF, MR, RS, TW

Resources: RD, KK, MD, JW, TW

Supervision: AP, RD, KK, MD, JW, TW

Validation: RD, KK, MD, JW, TW

Visualization: AP,TW

Writing – original draft: TW

Writing – review & editing: All authors.

## Competing interests

Authors declare that they have no competing interests.

## Data and materials availability

All data are available in the main text or the supplementary materials.

## Notes

### Competing Interest Statement

The authors have declared no competing interest.

## References

1. S. Mesquita F, Abrami L, Linder ME, Bamji SX, Dickinson BC, van der Goot FG. Mechanisms and functions of protein S-acylation. Vol. 25, Nature Reviews Molecular Cell Biology. Nature Research; 2024. p. 488–509.

2. Yang X, Chatterjee V, Ma Y, Zheng E, Yuan SY. Protein Palmitoylation in Leukocyte Signaling and Function. Vol. 8, Frontiers in Cell and Developmental Biology. Frontiers Media S.A.; 2020.

3. Greaves J, Chamberlain LH. Palmitoylation-dependent protein sorting. Journal of Cell Biology. 2007;176(3):249–54.

4. Buszka A, Pytyś A, Colvin D, Włodarczyk J, Wójtowicz T. S-Palmitoylation of Synaptic Proteins in Neuronal Plasticity in Normal and Pathological Brains. Vol. 12, Cells. MDPI; 2023.

5. Peng J, Liang D, Zhang Z. Palmitoylation of synaptic proteins: roles in functional regulation and pathogenesis of neurodegenerative diseases. Vol. 29, Cellular and Molecular Biology Letters. BioMed Central Ltd; 2024.

6. Zaręba-Kozioł M, Figiel I, Bartkowiak-Kaczmarek A, Włodarczyk J. Insights into protein S-palmitoylation in synaptic plasticity and neurological disorders: Potential and limitations of methods for detection and analysis. Vol. 11, Frontiers in Molecular Neuroscience. Frontiers Media S.A.; 2018.

7. Wlodarczyk J, Bhattacharyya R, Dore K, Ho GPH, Martin DDO, Mejias R, et al. Altered Protein Palmitoylation as Disease Mechanism in Neurodegenerative Disorders. Vol. 44, The Journal of neuroscienceL: the official journal of the Society for Neuroscience. 2024.

8. Globa AK, Bamji SX. Protein palmitoylation in the development and plasticity of neuronal connections. Vol. 45, Current Opinion in Neurobiology. 2017.

9. Tong J, Gao J, Feng B, Zhao X, Li J, Qi Y, et al. PPT1 Deciency-Induced GABAAR Hyperpalmitoylation Impairs Synaptic Transmission and Memory Formation. 2022; Available from: 10.21203/rs.3.rs-860958/v2

10. Kim SJ, Zhang Z, Sarkar C, Tsai PC, Lee YC, Dye L, et al. Palmitoyl protein thioesterase-1 deficiency impairs synaptic vesicle recycling at nerve terminals, contributing to neuropathology in humans and mice. Journal of Clinical Investigation. 2008 Sep 2;118(9):3075–86.

11. Li MD, Huang DH, Zheng YQ, Tian D, Ouyang H, Huang Z, et al. DHHC2-Mediated AKAP150 Palmitoylation Regulates Hippocampal Synaptic Plasticity and Fear Memory. 2022; Available from: 10.21203/rs.3.rs-2180782/v1

12. Li Y, Hu J, Höfer K, Wong AMS, Cooper JD, Birnbaum SG, et al. DHHC5 interacts with PDZ domain 3 of post-synaptic density-95 (PSD-95) protein and plays a role in learning and memory. Journal of Biological Chemistry. 2010 Apr 23;285(17):13022–31.

13. Kouskou M, Thomson DM, Brett RR, Wheeler L, Tate RJ, Pratt JA, et al. Disruption of the Zdhhc9 intellectual disability gene leads to behavioural abnormalities in a mouse model. Exp Neurol. 2018 Oct 1;308:35–46.

14. Urrego-Morales O, Gil-Lievana E, Ramirez-Mejia G, Francisco Rodríguez-Durán L, Lilia Escobar M, Delint-Ramirez I, et al. Inhibition of hippocampal palmitoyl acyltransferase activity impairs spatial learning and memory consolidation. Neurobiol Learn Mem. 2023 Apr 1;200.

15. Nasseri GG, Matin N, Wild AR, Tosefsky K, Flibotte S, Stacey RG, et al. Synaptic activity-dependent changes in the hippocampal palmitoylome. Sci Signal. 2022;15(763):eadd2519.

16. Malenka RC, Bear MF. LTP and LTD: An embarrassment of riches. Neuron. 2004;44(1):5–21.

17. Nicoll RA. A Brief History of Long-Term Potentiation. Neuron [Internet]. 2017;93(2):281–90. Available from: 10.1016/j.neuron.2016.12.015

18. Whitlock JR, Heynen AJ, Shuler MG, Bear MF. Learning induces long-term potentiation in the hippocampus. Science (1979). 2006 Aug 25;313(5790):1093–7.

19. Lüscher C, Malenka RC. NMDA receptor-dependent long-term potentiation and long-term depression (LTP/LTD). Cold Spring Harb Perspect Biol. 2012 Jun;4(6):1–15.

20. Baltaci SB, Mogulkoc R, Baltaci AK. Molecular Mechanisms of Early and Late LTP. Neurochem Res [Internet]. 2019;44(2):281–96. Available from: 10.1007/s11064-018-2695-4

21. Johnstone A, Mobley W. Local TrkB signaling: themes in development and neural plasticity. Available from: 10.1007/s00441-020-03278-7

22. Otmakhov N, Khibnik L, Otmakhova N, Carpenter S, Riahi S, Asrican B, et al. Forskolin-Induced LTP in the CA1 Hippocampal Region Is NMDA Receptor Dependent. J Neurophysiol. 2004 May;91(5):1955–62.

23. Szepesi Z, Hosy E, Ruszczycki B, Bijata M, Pyskaty M, Bikbaev A, et al. Synaptically released matrix metalloproteinase activity in control of structural plasticity and the cell surface distribution of GluA1-AMPA receptors. PLoS One. 2014 May 22;9(5).

24. Ojea Ramos S, Feld M, Fustiñana MS. Contributions of extracellular-signal regulated kinase 1/2 activity to the memory trace. Vol. 15, Frontiers in Molecular Neuroscience. Frontiers Media S.A.; 2022.

25. Brigidi GS, Bamji SX. Detection of protein palmitoylation in cultured hippocampal neurons by immunoprecipitation and acyl-biotin exchange (ABE). J Vis Exp. 2013;(72).

26. Kang R, Wan J, Arstikaitis P, Takahashi H, Huang K, Bailey AO, et al. Neural palmitoyl-proteomics reveals dynamic synaptic palmitoylation. Vol. 456, Nature. Nature Publishing Group; 2008. p. 904–9.

27. Brigidi GS, Santyr B, Shimell J, Jovellar B, Bamji SX. Activity-regulated trafficking of the palmitoyl-acyl transferase DHHC5. Nat Commun. 2015 Sep 3;6.

28. Xia ZX, Shen ZC, Zhang SQ, Wang J, Nie TL, Deng Q, et al. De-palmitoylation by N-(tert-Butyl) hydroxylamine inhibits AMPAR-mediated synaptic transmission via affecting receptor distribution in postsynaptic densities. CNS Neurosci Ther. 2019 Feb 1;25(2):187–99.

29. Webb Y, Hermida-Matsumoto L, Resh MD. Inhibition of Protein Palmitoylation, Raft Localization, and T Cell Signaling by 2-Bromopalmitate and Polyunsaturated Fatty Acids* [Internet]. 2000. Available from: http://www.jbc.org/

30. Zareba-Koziol M, Bartkowiak-Kaczmarek A, Figiel I, Krzystyniak A, Wojtowicz T, Bijata M, et al. Stress-induced Changes in the S-palmitoylation and S-nitrosylation of Synaptic Proteins. Molecular and Cellular Proteomics. 2019;18(10).

31. Chen RQ, Wang SH, Yao W, Wang JJ, Ji F, Yan JZ, et al. Role of glycine receptors in glycine-induced LTD in hippocampal CA1 pyramidal neurons. Neuropsychopharmacology. 2011 Aug;36(9):1948–58.

32. Brzdak P, Wójcicka O, Zareba-Koziol M, Minge D, Henneberger C, Wlodarczyk J, et al. Synaptic Potentiation at Basal and Apical Dendrites of Hippocampal Pyramidal Neurons Involves Activation of a Distinct Set of Extracellular and Intracellular Molecular Cues. Cerebral Cortex. 2019;29(1).

33. Nicoll RA, Schmitz D. Synaptic plasticity at hippocampal mossy fibre synapses. Nat Rev Neurosci. 2005;6(11):863–76.

34. Yilmaz-Rastoder E, Miyamae T, Braun AE, Thiels E. LTP- and LTD-inducing stimulations cause opposite changes in arc/arg3.1 mRNA level in hippocampal area CA1 in vivo. Hippocampus. 2011 Dec;21(12):1290– 301.

35. Dziembowska M, Milek J, Janusz A, Rejmak E, Romanowska E, Gorkiewicz T, et al. Activity-dependent local translation of matrix metalloproteinase-9. Journal of Neuroscience. 2012 Oct 17;32(42):14538–47.

36. Corera AT, Doucet G, Fon EA. Long-term potentiation in isolated dendritic spines. PLoS One. 2009 Jun 23;4(6).

37. Li MD, Wang L, Zheng YQ, Huang DH, Xia ZX, Liu JM, et al. DHHC2 regulates fear memory formation, LTP, and AKAP150 signaling in the hippocampus. iScience. 2023 Sep 15;26(9).

38. Valtorta F, Pennuto M, Bonanomi D, Benfenati F. Synaptophysin: Leading actor or walk-on role in synaptic vesicle exocytosis? Vol. 26, BioEssays. 2004. p. 445–53.

39. Granseth B, Odermatt B, Royle SJ, Lagnado L. Clathrin-mediated endocytosis: The physiological mechanism of vesicle retrieval at hippocampal synapses. In: Journal of Physiology. 2007. p. 681–6.

40. Pyeon HJ, Lee YI. Differential expression levels of synaptophysin through developmental stages in hippocampal region of mouse brain. Anat Cell Biol. 2012;45(2):97.

41. Kokotos AC, Harper CB, Marland JRK, Smillie KJ, Cousin MA, Gordon SL. Synaptophysin sustains presynaptic performance by preserving vesicular synaptobrevin-II levels. J Neurochem. 2019 Oct 1;151(1):28–37.

42. Bacci A, Coco S, Pravettoni E, Schenk U, Armano S, Frassoni C, et al. Chronic Blockade of Glutamate Receptors Enhances Presynaptic Release and Downregulates the Interaction between Synaptophysin-Synaptobrevin-Vesicle-Associated Membrane Protein 2. 2001.

43. mullany1998.

44. Kraev I V, Chaudhury S, Davies HA, Dallérac G, Doyère V, Stewart MG. Long Term Potentiation (LTP) and Long Term Depression (LTD) Cause Differential Spatial Redistribution of the Synaptic Vesicle Protein Synaptophysin in the Middle Molecular Layer of the Dentate Gyrus in Rat Hippocampus. Opera Med Physiol. 2016;2(4):205.

45. Regehr WG. Short-term presynaptic plasticity. Cold Spring Harb Perspect Biol. 2012 Jul;4(7):1–19.

46. Shen ZC, Xia ZX, Liu JM, Zheng JY, Luo YF, Yang H, et al. APT1-Mediated Depalmitoylation Regulates Hippocampal Synaptic Plasticity. Journal of Neuroscience. 2022 Mar 30;42(13):2662–77.

47. Shinozaki K, Kume H, Kuzume H, Obata K, Maruyama K. Interactive report Norbin, a neurite-outgrowth-related protein, is a cytosolic protein localized in the somatodendritic region of neurons and distributed prominently in dendritic outgrowth in Purkinje cells 1 [Internet]. Vol. 71, Molecular Brain Research. 1999. Available from: www.elsevier.comrlocaterbres

48. Chetwynd SA, Andrews S, Inglesfield S, Delon C, Ktistakis NT, Welch HCE. Functions and mechanisms of the GPCR adaptor protein Norbin. Vol. 51, Biochemical Society Transactions. Portland Press Ltd; 2023. p. 1545–58.

49. Westin L, Reuss M, Lindskog M, Aperia A, Brismar H. Nanoscopic spine localization of Norbin, an mGluR5 accessory protein [Internet]. Vol. 15, BMC Neuroscience. 2014. Available from: http://www.biomedcentral.com/1471-2202/15/45

50. Fatima A, Hoeber J, Schuster J, Koshimizu E, Maya-Gonzalez C, Keren B, et al. Monoallelic and bi-allelic variants in NCDN cause neurodevelopmental delay, intellectual disability, and epilepsy. Am J Hum Genet. 2021 Apr 1;108(4):739–48.

51. Dateki M, Horii T, Kasuya Y, Mochizuki R, Nagao Y, Ishida J, et al. Neurochondrin negatively regulates CaMKII phosphorylation, and nervous system-specific gene disruption results in epileptic seizure. Journal of Biological Chemistry. 2005 May 27;280(21):20503–8.

52. Oku S, Takahashi N, Fukata Y, Fukata M. In Silico screening for palmitoyl substrates reveals a role for DHHC1/3/10 (zDHHC1/3/11)-mediated neurochondrin palmitoylation in its targeting to Rab5-positive endosomes. Journal of Biological Chemistry. 2013 Jul 5;288(27):19816–26.

53. Schwaibold EMC, Brandt DT. Identification of Neurochondrin as a new interaction partner of the FH3 domain of the Diaphanous-related formin Dia1. Biochem Biophys Res Commun. 2008 Aug 29;373(3):366– 72.

54. Dityatev A, Bukalo O, Schachner M. Modulation of synaptic transmission and plasticity by cell adhesion and repulsion molecules. Vol. 4, Neuron Glia Biology. 2008. p. 197–209.

55. Niethammer P, Delling M, Sytnyk V, Dityatev A, Fukami K, Schachner M. Cosignaling of NCAM via lipid rafts and the FGF receptor is required for neuritogenesis. Journal of Cell Biology. 2002 Apr 29;157(3):521– 32.

56. Ponimaskin E, Dityateva G, Ruonala MO, Fukata M, Fukata Y, Kobe F, et al. Fibroblast growth factor-regulated palmitoylation of the neural cell adhesion molecule determines neuronal morphogenesis. Journal of Neuroscience. 2008 Sep 3;28(36):8897–907.

57. Diestel S, Schaefer D, Cremer H, Schmitz B. NCAM is ubiquitylated, endocytosed and recycled in neurons. J Cell Sci. 2007 Nov 15;120(22):4035–49.

58. Lüthi A, Laurent JP, Figurovt A, Mullert D, Schachnert M. Hippocampal long-term potentiation and neural cell adhesion molecules L1 and NCAM. Nature [Internet]. 1994;372(6508):777–9. Available from: 10.1038/372777a0

59. Ohyama T, Verstreken P, Ly C V., Rosenmund T, Rajan A, Tien AC, et al. Huntingtin-interacting protein 14, a palmitoyl transferase required for exocytosis and targeting of CSP to synaptic vesicles. Journal of Cell Biology. 2007 Dec 31;179(7):1481–96.

60. Dejanovic B, Semtner M, Ebert S, Lamkemeyer T, Neuser F, Lüscher B, et al. Palmitoylation of Gephyrin Controls Receptor Clustering and Plasticity of GABAergic Synapses. PLoS Biol. 2014;12(7):1–16.

61. Valenzuela A, Meservey L, Nguyen H, Fu M meng. Golgi Outposts Nucleate Microtubules in Cells with Specialized Shapes. Vol. 30, Trends in Cell Biology. Elsevier Ltd; 2020. p. 792–804.

62. Chojnacka M, Beroun A, Magnowska M, Stawikowska A, Cysewski D, Milek J, et al. Impaired synaptic incorporation of AMPA receptors in a mouse model of fragile X syndrome. Front Mol Neurosci. 2023;16.

63. Gorenberg EL, Tieze SM, Yücel B, Zhao HR, Chou V, Wirak GS, et al. Identification of substrates of palmitoyl protein thioesterase 1 highlights roles of depalmitoylation in disulfide bond formation and synaptic function. PLoS Biol. 2022 Mar 1;20(3).

64. El-Husseini AED, Schnell E, Dakoji S, Sweeney N, Zhou Q, Prange O, et al. Synaptic strength regulated by palmitate cycling on PSD-95. Cell. 2002 Mar 22;108(6):849–63.

65. Kleene R, Mzoughi M, Joshi G, Kalus I, Bormann U, Schulze C, et al. NCAM-induced neurite outgrowth depends on binding of calmodulin to NCAM and on nuclear import of NCAM and fak fragments. Journal of Neuroscience. 2010 Aug 11;30(32):10784–98.

66. Boxer AL, Moreno H, Rudy B, Ziff EB. FGF-2 Potentiates Ca 2-Dependent Inactivation of NMDA Receptor Currents in Hippocampal Neurons. 1999.

67. Milnerwood AJ, Parsons MP, Young FB, Singaraja RR, Franciosi S, Volta M, et al. Memory and synaptic deficits in Hip14/DHHC17 knockout mice. Proc Natl Acad Sci U S A. 2013 Dec 10;110(50):20296–301.

68. Sapir T, Segal M, Grigoryan G, Hansson KM, James P, Segal M, et al. The interactome of palmitoyl-protein thioesterase 1 (PPT1) affects neuronal morphology and function. Front Cell Neurosci. 2019 Jan 29;13.

69. Spencer-Segal JL, Waters EM, Bath KG, Chao M V., McEwen BS, Milner TA. Distribution of phosphorylated Trkb receptor in the mouse hippocampal formation depends on sex and estrous cycle stage. Journal of Neuroscience. 2011 May 4;31(18):6780–90.

70. Navakkode S, Sajikumar S, Korte M, Soong TW. Dopamine induces LTP differentially in apical and basal dendrites through BDNF and voltage-dependent calcium channels. Learning and Memory. 2012 Jul;19(7):294–9.

71. Guskjolen A, Cembrowski MS. Engram neurons: Encoding, consolidation, retrieval, and forgetting of memory. Vol. 28, Molecular Psychiatry. Springer Nature; 2023. p. 3207–19.

72. Wójtowicz T, Brzdąk P, Mozrzymas JW. Diverse impact of acute and long-term extracellular proteolytic activity on plasticity of neuronal excitability. Vol. 9, Frontiers in Cellular Neuroscience. Frontiers Media S.A.; 2015.

73. Chiappalone M, Massobrio P, Martinoia S. Network plasticity in cortical assemblies. European Journal of Neuroscience. 2008 Jul;28(1):221–37.

74. Shen ZC, Liu JM, Zheng JY, Li MD, Tian D, Pan Y, et al. Regulation of anxiety-like behaviors by S-palmitoylation and S-nitrosylation in basolateral amygdala. Biomedicine and Pharmacotherapy. 2023 Dec 31;169.

75. Michaluk P, Wawrzyniak M, Alot P, Szczot M, Wyrembek P, Mercik K, et al. Influence of matrix metalloproteinase MMP-9 on dendritic spine morphology. J Cell Sci. 2011 Oct 1;124(19):3369–80.

76. Kuzniewska B, Chojnacka M, Milek J, Dziembowska M. Preparation of polysomal fractions from mouse brain synaptoneurosomes and analysis of polysomal-bound mRNAs. J Neurosci Methods. 2018 Jan 1;293:226–33.

77. Chiappalone M, Massobrio P, Martinoia S. Network plasticity in cortical assemblies. European Journal of Neuroscience. 2008;28(1).

78. Schindelin J, Arganda-Carreras I, Frise E, Kaynig V, Longair M, Pietzsch T, et al. Fiji: An open-source platform for biological-image analysis. Vol. 9, Nature Methods. 2012. p. 676–82.

79. Myers SA, Rhoads A, Cocco AR, Peckner R, Haber AL, Schweitzer LD, et al. Streamlined protocol for deep proteomic profiling of FAC-sorted cells and its application to freshly isolated murine immune cells. Molecular and Cellular Proteomics. 2019 May 1;18(5):995–1009.

80. Singh N, Golime RR, Acharya J, Palit M. Quantitative proteomic changes after organophosphorous nerve agent exposure in the rat hippocampus. ACS Chem Neurosci. 2020 Sep 2;11(17):2638–48.

