## Supplementary figures for "Temporal and protein-specific S-palmitoylation supports synaptic and neural network plasticity"

Agata Pytyś *et al.*

**This PDF file includes:**

Figs. S1 to S5
Data S1

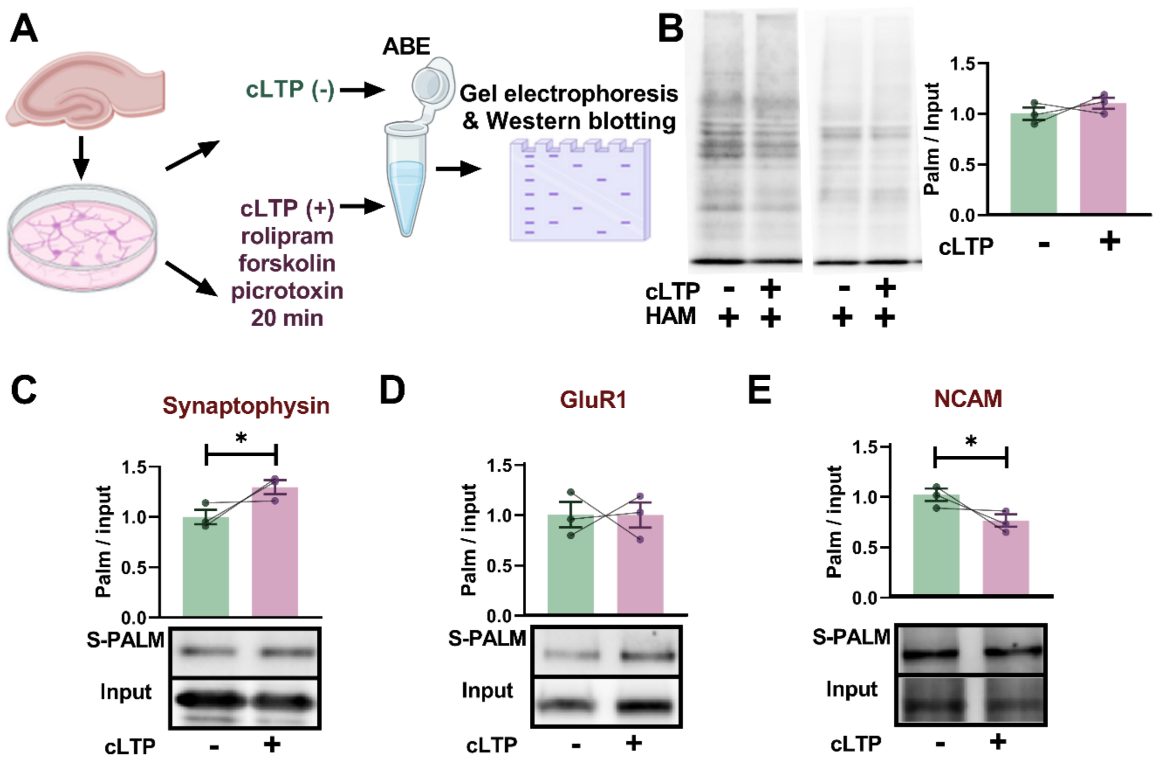


**Fig. S1.**

**Induction of synaptic plasticity in neuronal cultures regulates palmitoylation of synaptic proteins.** **(A)** Primary rat hippocampal neurons were cultured for 14 days and treated with cLTP cocktail containing rolipram, forskolin and picrotoxin (cLTP +) or solvent (cLTP -). Culture homogenates were collected 20 min after treatment, subjected to ABE and S-PALM and input fractions were immunoblotted for target proteins, as indicated. **(B)** Western blot of global palmitoylation 20 min post cLTP indicated no shift in global protein palmitoylation ($n$ = 3 cultures, $p$ = 0.27, unpaired Student's t-test). **(C)** Western blot and quantification of the ABE assay described in (A) performed on selected presynaptic proteins. The levels of palmitoylated synaptophysin increased significantly post cLTP ($n$ = 3 cultures, $p$ = 0.039, unpaired Student's t-test). **(D)** Western blot and quantification of the ABE assay described in (A) performed on selected postsynaptic proteins. **(E)** Western blot and quantification of the ABE assay described in (A) performed on exemplary bipolar adhesion molecule NCAM. The levels of palmitoylated NCAM increased significantly post cLTP ($n$ = 3 cultures, $p$ = 0.041, unpaired Student's t-test). Data are means ± SEM from $n$ = 3 to 5 animals. *$p < 0.05$.

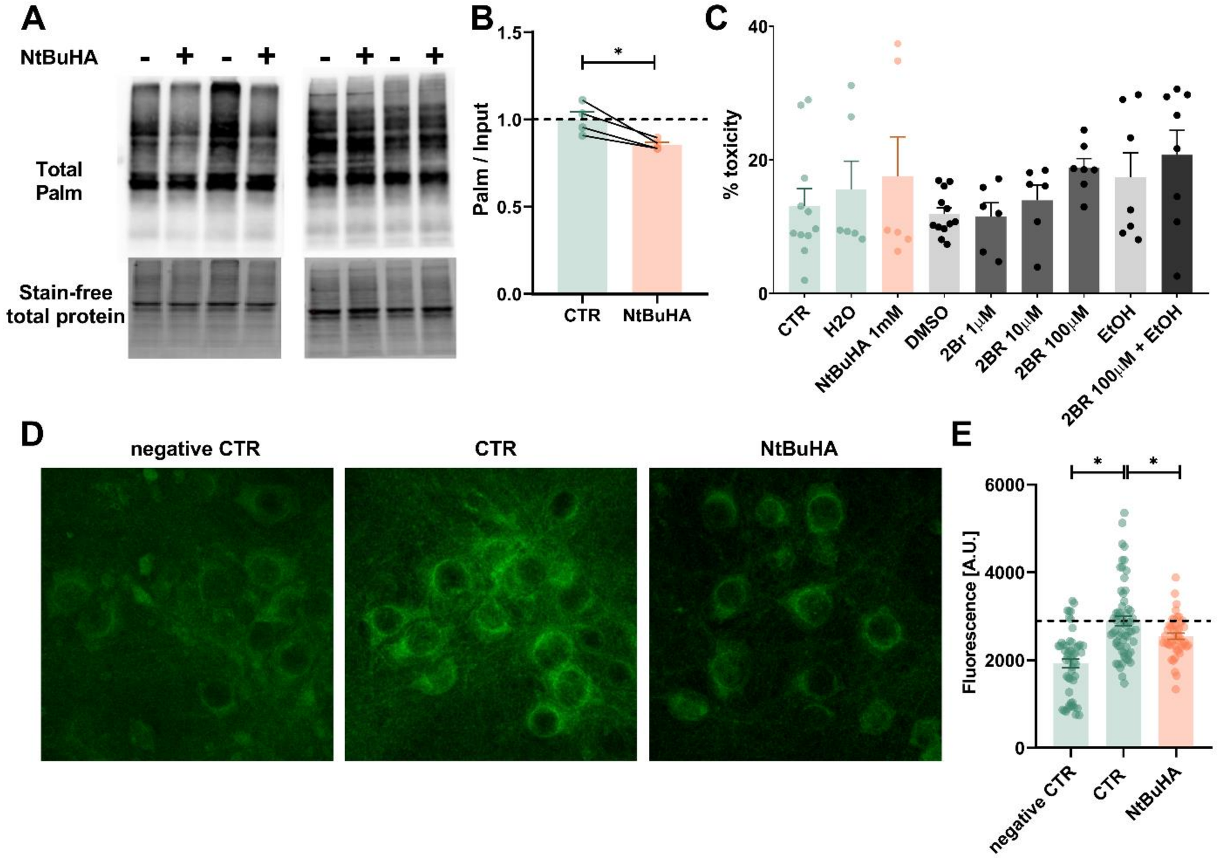


**Fig S2.**

**Deacylation of proteins in primary hippocampal cultures with NtBuHA.**

**(A)** Primary rat hippocampal cultures at 14 DIV were treated overnight with 1 mM NtBuHA. Global protein palmitoylation was assessed using HRP-streptavidin after the acyl-biotin exchange (ABE) protocol on whole-culture homogenates. **(B)** Quantification of the Western blot shown in (A). NtBuHA significantly reduced S-PALM of proteins when normalized to the loading control (stain-free total protein intensity; *n* = 4 cultures; *p* = 0.021, unpaired Student's *t*-test). **(C)** Primary hippocampal cultures at 14 DIV were treated overnight with 1 mM NtBuHA (dissolved in water) or with 2BR (1–100 μM) dissolved in DMSO (0.1%) or ethanol (EtOH, 0.1%). Culture media were collected and analyzed for lactate dehydrogenase (LDH) release using the CytoTox 96 Cytotoxicity Assay. Only 2-BR at 100 μM dissolved in DMSO caused a significant increase in LDH release, indicating elevated cell mortality ($n \geq 6$ cultures; one-way ANOVA, $F$ (7, 56) = 2.185, $p$ = 0.049). **(D)** Exemplary images of total proteome palmitoylation in neuronal cultures subjected to click chemistry reaction with Oregon Green 488 dye. Primary rat hippocampal cultures at 14 DIV were treated overnight with (CTR) or without exogenous alkine palmitate (negative CTR). Additionally, 1 mM NtBuHA was added to some cultures. **(E)** Quantification of average fluorescence intensity of Oregon Green in cultures described in (D). Control cultures had significantly larger fluorescence compared to negative CTR and NtBuHA treated cultures ($n \geq 45$ pictures per group, $N = 2$ cultures, $F$ (2, 156) = 25.69, $p < 0.001$, one-way ANOVA with Tukey post-hoc). Data are means ± SEM. Asterisks indicate statistical significance: * $p < 0.05$.

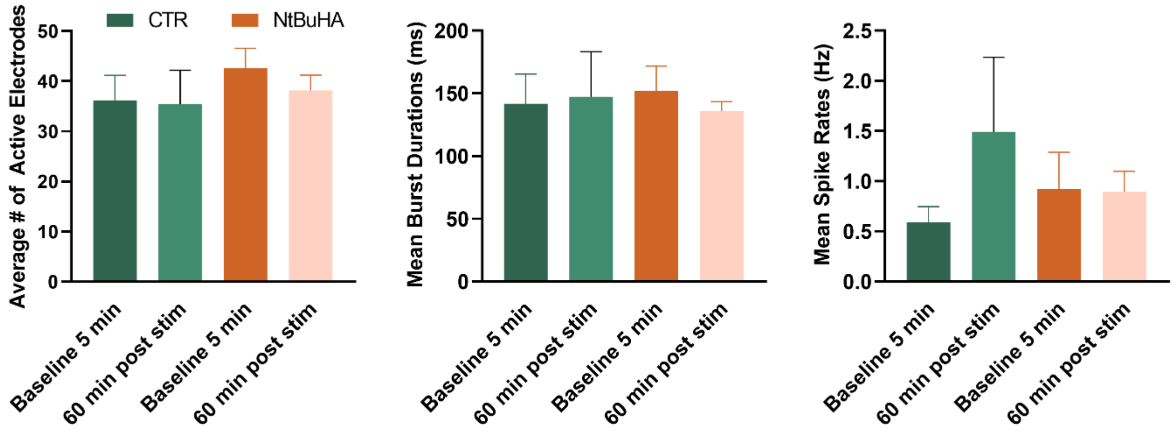


**Fig S3.**

**Protein deacylation does not affect neuronal spiking *in vitro*.**

Primary rat hippocampal neurons were cultured for 14 days on multielectrode arrays (MEAs), enabling recordings of action potentials from up to 60 network sites. Spikes were quantified in control cultures (green) and cultures treated overnight with NtBuHA (orange) before and after inducing network plasticity through associative activity (stim.). There were no significant main effects of the drug or stimulation on the number of active electrodes, burst duration, or mean spike firing rate ($F(1,8) = 0.135$, $p = 0.71$; $F(1,10) = 0.24$, $p = 0.63$; $F(1,16) = 1,15$, $p = 0.29$, two-way ANOVA, treatment x stimulation). Data are shown as mean ± SEM, with $n$ = 5 MEAs per group.

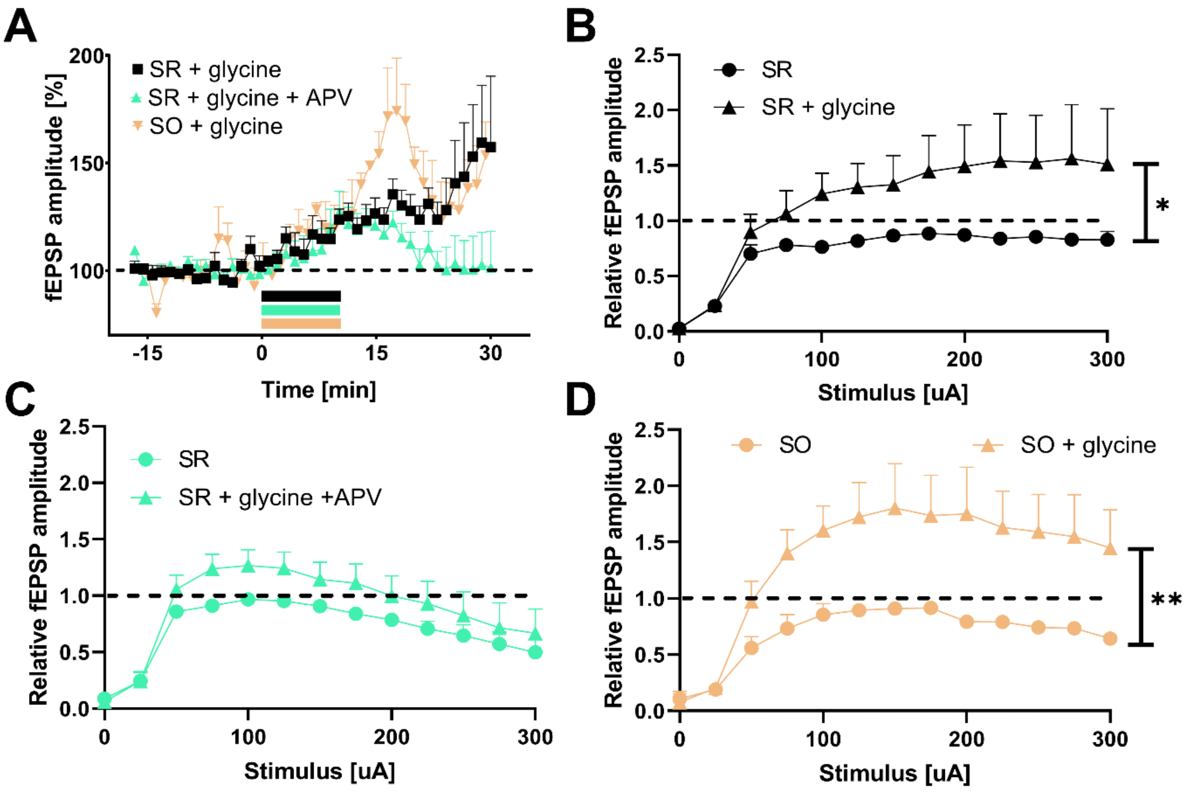


**Fig S4.**

**Glycine-induced NMDAR-dependent synaptic potentiation in hippocampal slices.**

**(A)** Time course of fEPSPs recorded in stratum radiatum (SR) or stratum oriens (SO) of CA1 hippocampal region in response to extracellular stimulation of Schaffer collaterals. Addition of glycine (600µM, at time 0 min. for 10 minutes) in $Mg^{2+}$-free aCSF resulted in an enhancement of fEPSP amplitudes both in SR and SO. In SR this effect was not observed when glycine was applied in the presence of NMDAR antagonist APV (50µM) (n=7-8 slices per group). (B-D) Statistical analysis of fEPSP amplitudes recorded in response to a wide range of stimuli before and 30 minutes after application of glycine in slices shown in A. **(B)** Glycine significantly enhanced fEPSP amplitudes ($n$ = 8 slices per group, $F(12, 168) = 1.831$, $p$ = 0.04, two-way ANOVA, treatment × stimulus intensity). This effect was not observed in the presence of APV as shown in **(C)** ($n$ = 7 slices per group, $F(12, 144) = 0.86$, $p = 0.58$, two-way ANOVA, treatment × stimulus intensity). **(D)** Glycine significantly enhanced fEPSP amplitudes in SO ($n = 4$ slices per group, $F(12, 72) = 2.947$, $p = 0.002$, two-way ANOVA, treatment × stimulus intensity). Asterisks indicate statistical significance: * $p < 0.05$, ** $p < 0.01$.

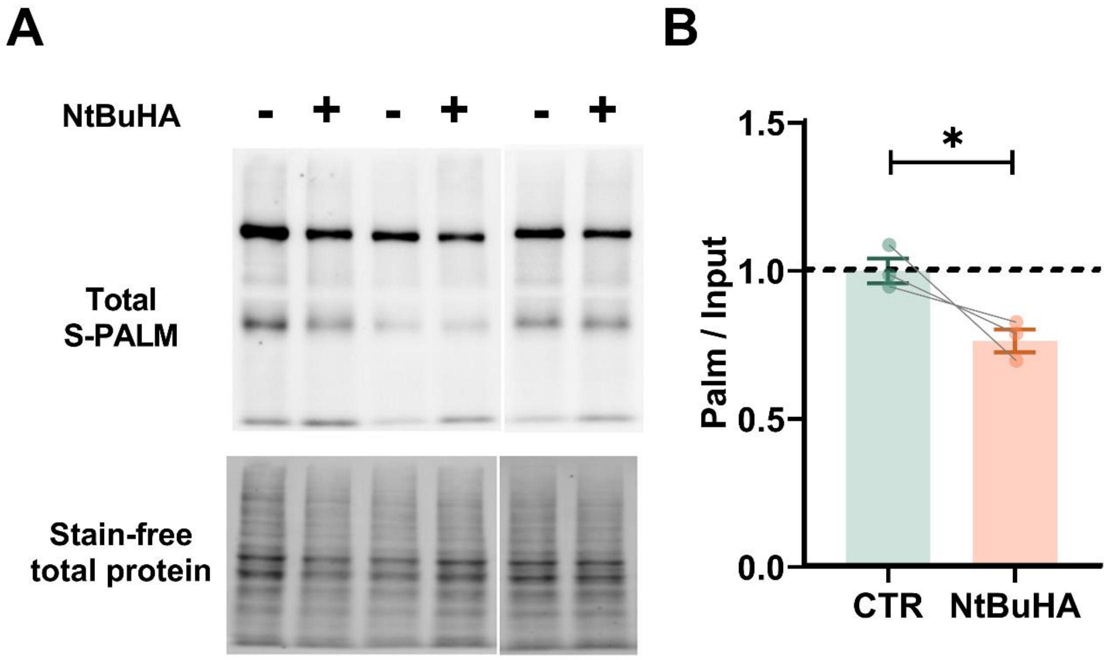


**Fig S5.**

**Deacylation of proteins in hippocampal slices with NtBuHA.**

**(A)** Hippocampi from Wistar rats aged 45-60 days were cut in slices (350µm thick) and left in either aCSF (CTR) or aCSF containing NtBuHA (1mM) for 1-4 hours. Global protein palmitoylation was assessed using HRP-streptavidin after the acyl-biotin exchange (ABE) protocol on tissue homogenates. **(B)** Quantification of WB shown in (A). NtBuHA significantly reduced S-palmitoylation of proteins when normalized to the loading control (stain-free total protein intensity; $n$ = 3 animals; $p = 0.014$, unpaired Student's $t$-test). Asterisks indicate statistical significance: * $p < 0.05$.

**Data S1.**

**Palmitoyl-proteins from this proteomic study of synaptoneurosomes.**

The Excel spreadsheet 1 contains the list of all input proteins. Sheet 2 shows the list of all palmitoylated proteins. Sheet 3 shows differentially palmitoylated proteins detected in 37ºC and STIM 37ºC samples. Sheets 4-7 show results of KEGG and GO functional protein enrichment analyses for palmitoylated proteins described in Sheet 3 upon comparison to all proteins reported in homogenates of rat hippocampus shown in Sheet 8 (1).
